# An experimentally tractable framework for quantitative and integrative analysis of aggregation in the origin of animals

**DOI:** 10.1101/2025.05.14.653760

**Authors:** Gonzalo Bercedo-Saborido, Daria Stepanova, Iñaki Ruiz-Trillo, Tomàs Alarcón, Koryu Kin

**Author notes:** These authors contributed equally to this work (Gonzalo Bercedo-Saborido, Daria Stepanova).

## Abstract

The origin of animals from their unicellular ancestors remains a fundamental biological question. Cell aggregation, a widespread eukaryotic behaviour, has been underappreciated as a potential pathway to multicellularity. Nevertheless, aggregation is a key mechanism present in extant animals and has been described in both choanoflagellates and filastereans, the closest unicellular relatives of animals. Sebé-Pedrós et al. (2013) established that aggregative, filopodial and cystic cultures of *Capsaspora owczarzaki* have distinct transcriptomes. Here we extend this description by following serum-induced aggregation and disaggregation over 72 h using live imaging and RNA-seq at eight sampling times. Comparison with reanalysed 2013 data reveals changing correspondence to the established life-stage programs. The nominal 3-min sample shows an immediate RNA-abundance response distinct from the aggregative-versus-filopodial contrast. Aggregative-associated genes are enriched from 12 h onwards, while cystic-associated genes become enriched at 60-72 h alongside continued aggregative enrichment. Homologues of animal adhesion and signalling genes show dynamic expression. In parallel, we developed a novel quantitative mathematical model, calibrated with experimental data, that captures the full aggregation cycle, and identified a nonlinear threshold-like adhesion response to fetal bovine serum (chemical inducer) coupled to its local depletion, sufficient to explain and predict aggregate formation, growth, and disassembly. Together, the transcriptomic data and the mathematical model resolve temporal features missed by endpoint sampling in the previous study and provide an integrated experimental-theoretical framework to interrogate how facultative aggregation could have influenced the emergence of animal multicellularity.

## Introduction

How animals evolved from their unicellular ancestor remains a major biological question. Comparative genomics and cell biology have shown that the unicellular relatives of animals already possessed many genes later used in animal multicellularity (Richter et al., 2018; Ros-Rocher et al., 2021; Ruiz-Trillo et al., 2023). However, these discoveries have not resolved the cellular context in which such genes originally functioned, nor which cellular route first stabilized them into multicellular organization. Current reconstructions therefore remain open to multiple possibilities, including clonal development, multinucleate (coenocyte) states, and facultative aggregation (Ruiz-Trillo et al., 2023).

Among these possibilities, aggregation has classically been considered an unlikely route due to the potential negative consequences of the emergence of cheaters (Grosberg & Strathmann, 2007; Michod & Roze, 2001; Pentz et al., 2023). Yet this concern does not by itself rule aggregation out as a relevant evolutionary route (La Fortezza & Velicer, 2021; Ostrowski, 2019; Pande & Velicer, 2018). Notably, cell aggregation is indeed a powerful way to become multicellular. First, cell aggregation is widespread in eukaryotes including animals (Brunet & King, 2017; Chaine et al., 2010; Du et al., 2015; Kawabe et al., 2019; Kidner et al., 2024; Kin & Schaap, 2021; Sebé-Pedrós et al., 2017); second, aggregation can provide immediate ecological advantages, such as rapid predator avoidance and improved extracellular metabolism (Herron et al., 2019; Pentz et al., 2020; Tong et al., 2022); third, in extant animals, collective cell behaviours play important roles in development, tissue repair, and homeostasis (Curtis, 1962; Friedl & Gilmour, 2009; Hall & Miyake, 2000; Park et al., 2017; Ruperti et al., 2024). Moreover, because aggregating cells can have more free movement than in clonal colonies or multinucleate structures, aggregation may provide a distinct route by which collective organization, cell sorting and transient division of labour could emerge (Newman, 2020; Newman et al., 2006; Ruiz-Trillo et al., 2023).

To evaluate whether aggregation could have contributed to the origin of animal multicellularity, we need to know in unicellular relatives of animals whether aggregation is an actively regulated biological process, if it proceeds through reproducible temporal states, and whether genes relevant to animal multicellularity are dynamically deployed during those states. If cell adhesion, signalling, cytoskeletal, or regulatory genes shared with animals are activated in a coordinated fashion during aggregation, this would suggest that aggregative multicellularity may have provided an ancestral context in which such genes were used, refined, or later co-opted into more stable multicellular systems.

Recent work has strengthened the relevance of aggregation to animal origins. In particular, aggregative multicellularity has now been described in multiple close unicellular relatives of animals, including choanoflagellates and filastereans (Li et al., 2026; Ros-Rocher et al., 2026). A new study of the filasterean *Ministeria vibrans* (Li et al., 2026) showed that aggregation is indeed accompanied by the deployment of homologs of animal genes involved in adhesion, signalling and transcriptional regulation. These findings suggest that the use of animal multicellularity genes in aggregation is not restricted to a single species or group and may reflect a broader pre-metazoan cellular context in which such genes were used before being co-opted into animal development.

The filasterean *Capsaspora owczarzaki* (hereafter *Capsaspora,* Figure 1) is well suited for studying the role of aggregation in the premetazoan context. As one of the closest known relatives of animals, its genome shares a large set of “multicellular” genes with animals, including a complete integrin adhesome, the Hippo pathway, several protein tyrosine kinases, and various developmental transcription factors (TFs) (Suga et al., 2013). Importantly, *Capsaspora* can be cultured axenically (in contrast to *M. vibrans* or choanoflagellates) minimizing confounding microbial interactions, its aggregation can be reliably induced by specific chemical triggers (components of FBS) (Ros-Rocher et al., 2023), and emerging transfection and genome-editing tools make it a promising system for future causal perturbation (Phillips et al., 2022).

**Figure 1.**
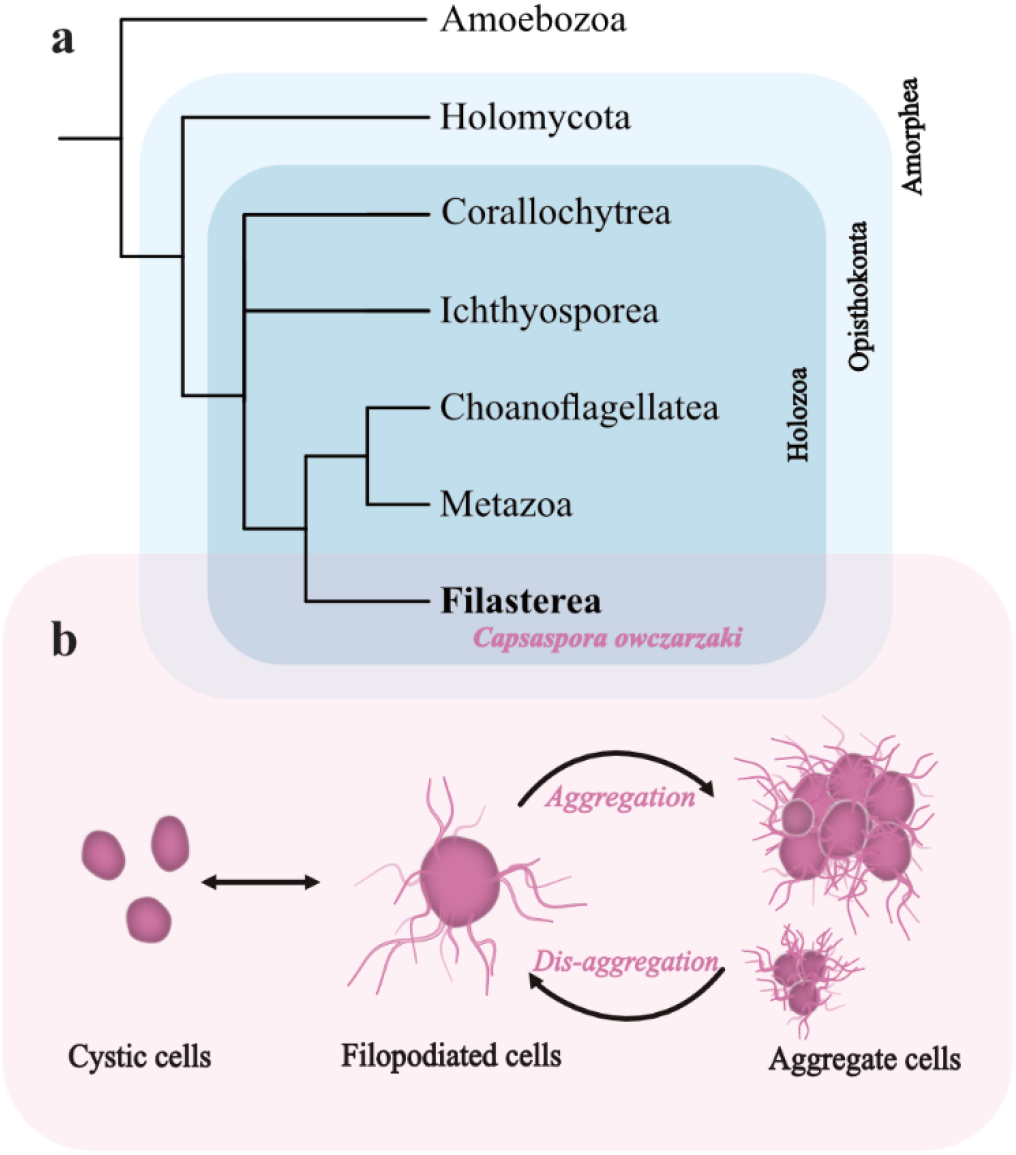
Unicellular holozoans and *Capsaspora.* **a**, Phylogenetic relationship of the unicellular closest relatives of animals based on (Ocaña-Pallarès et al., 2022; Ros-Rocher et al., 2021). The filasterean *Capsaspora owczarzaki* is among the best studied relatives of animals, and it shares with animals many genes that were thought to be animal-specific (Suga et al., 2013). b, *Capsaspora* presents three different life stages under culture conditions (Sebé-Pedrós et al., 2013). Filopodial stage in which single amoeboid cells adhere to the substrate. Under maintained stressful conditions cells can enter a cystic stage losing their filopodia and forming resistant structures; this stage is reversible upon recovering optimal conditions. *Capsaspora* cells can also form a multicellular stage by aggregation, in which cells form larger clusters through filopodial interconnections and formation of an extracellular matrix, exhibiting a high plasticity regarding their size.

In *Capsaspora*, Sebé-Pedrós et al. described distinct life-stage transcriptomes, including increased expression of integrin-adhesome components and tyrosine kinases in mature aggregates induced by orbital agitation (Sebé-Pedrós et al., 2013). However, that endpoint comparison did not resolve the timing of expression changes during induction, growth and disassembly. Temporal transcriptomics can address this gap, while a quantitative model can test whether specified cell-level assumptions reproduce observed collective dynamics. These approaches offer complementary descriptions; neither expression correlations nor successful model fits alone identify a molecular mechanism.

Here we extend the life-stage analysis of Sebé-Pedrós et al. (2013) with live imaging and an eight-point RNA-seq time course of FBS-induced aggregation and disaggregation. We compare within-study expression changes using a common annotation to determine how the earlier life-stage programs relate to the time course. In parallel, we develop an agent-based model to explore whether FBS-dependent effective adhesion, motility, proliferation and local inducer depletion can reproduce the observed dynamics. Our morphological and genetic characterization of cell aggregation and disaggregation in *Capsaspora* demonstrates how genes involved in multicellular functions are dynamically expressed throughout these processes. Moreover, the mathematical model we constructed describes the kinetics of the aggregation process and shows that differential cell-cell adhesion and compactness are driving its dynamics. Overall, our results highlight a potential role of cell aggregation in discussions of animal origins and position *Capsaspora* as an integrative model system for quantitatively evaluating how aggregation may have paved the way for animal multicellularity.

## Results

### The aggregative behaviour of Capsaspora owczarzaki is highly reproducible, with sequential stages that can be identified throughout its temporal dynamics

The formation of *Capsaspora* aggregates has been previously described, including the chemical cues necessary for inducing the aggregation process (Ros-Rocher et al., 2023). There is also an RNA-seq analysis of the different life stages of *Capsaspora* (amoeboid, cystic, and aggregative), that shows different transcriptomic profiles for each life stage (Sebé-Pedrós et al., 2013). However, the previous analysis showed only a specific moment of aggregation in late mature aggregates induced by orbital agitation. If we want to understand the potential role of aggregation in animal origins, we need a systematic approach to the overall dynamics of cell aggregation to unravel which genes are dynamically employed, establishing a system in which genetic perturbations can be quantitatively assessed. We here specifically examined whether *Capsaspora’s* multicellular aggregative behaviour is deploying “multicellular genes” that later on were key for animal multicellularity. To this end, it was crucial to first identify key intermediate stages of the aggregation process. Microscopic observation guided the selection of eight time points of the aggregation process in which a clear change in aggregation dynamics was observed in a reproducible manner (Movie 1) (Figure 2).

**Figure 2.**
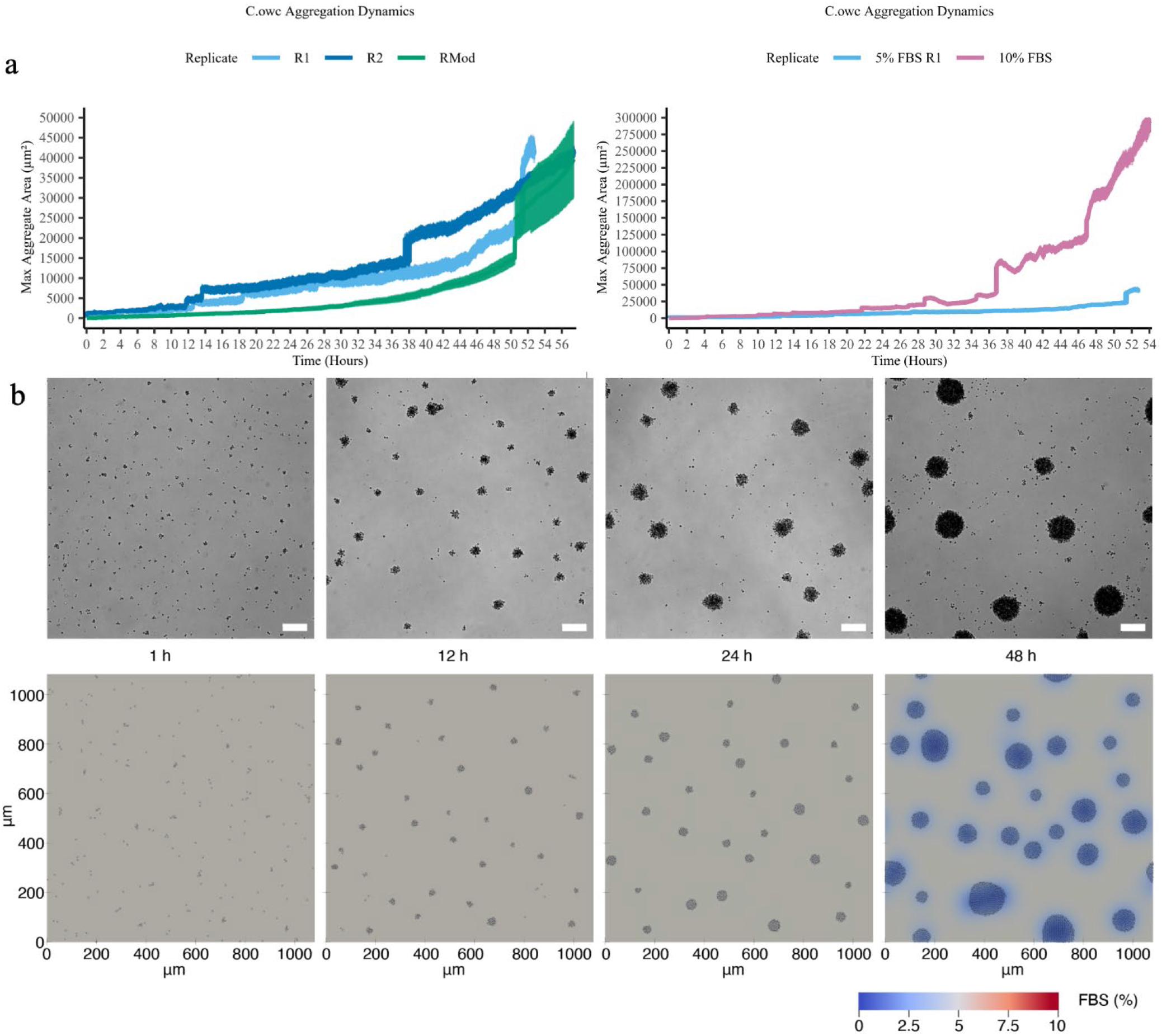
Comparison of *Capsaspora* aggregation dynamics between experimental results and model simulations under 5% FBS. **a**, *Comparison of Capsaspora aggregate growth dynamics* with different FBS concentrations. The left panel shows the aggregation dynamics of two experimental replicates at 5% FBS (R1 and R2) and a simulation of the mathematical model at 5% FBS (Rmod, green line). The right panel shows experimental results comparing the aggregation dynamics at 5% and 10% FBS. The growth and formation of larger aggregates occurs through two mechanisms, the first involves cell proliferation within aggregates, reflected by a continuous and gradual increase in aggregate area. The second occurs through the clustering of pre-existing aggregates, resulting in sharp increases in the aggregate area. **b,** Experimental images scale bar: 100 μm (top) vs Model snapshots (bottom) comparison, numerical simulation at 1, 12, 24 and 48 hours after administration of 5% FBS. Both experimental and simulated images cover a 1080 x 1080 μm domain. In the simulations, cell colours represent the local FBS concentration, which is also reflected in the background of the medium. The colour bar on the right indicates FBS levels: initial 5% FBS appears grey, while depletion due to metabolic activity of *Capsaspora* shifts colours to blue, with dark blue indicating no FBS. Numerical simulations were performed with sigmoidal Hill response of cell adhesion strength to FBS levels (see Supplementary Information) and all parameter values are as Extended in Data Table 1.

We start from a confluent culture of single cells seeded in media without fetal bovine serum (FBS) (T0). At time point 1 (T1, 3 min after aggregation induction), cells detach from the surface and begin to get close to one another, forming aggregates in groups of ∼5 cells. At time point 2 (T2, 12 h), smaller aggregates of approximately 20-30 cells are formed. By time point 3 (T3, 24 h), growth of the aggregates is mainly generated through cell division, with the number of cells typically reaching hundreds; this increase is represented by a smooth increase in the maximum area of the aggregates shown in the graph. At time point 4 (T4, 36 h), merging between aggregates becomes more frequent, leading to the formation of macroscopic aggregates. After time point 5 (T5, 48 h), the aggregates are reaching their maximum size, and by time point 6 (T6, 60 h), dissociation becomes visible from their centre. Finally, at time point 7 (T7, 72 h), the aggregates revert to single cells.

### Aggregation via FBS-upregulated adhesion

Experimental studies have shown that FBS components, such as lipoproteins and calcium ions, induce reversible aggregation in *Capsaspora*. When FBS is abundant, *Capsaspora* metabolises its components, forming aggregates that merge into larger clusters over time. Once aggregates are disaggregated into single cells, the re-addition of FBS can rapidly induce reaggregation of cells (Ros-Rocher et al., 2023). Motivated by these observations, we developed a mathematical model to capture this chemically induced aggregation process (see Extended Data Figure 1).

Our model integrates individual cell behaviour with local FBS concentration, which decreases over time due to consumption by cellular metabolism. Cells move within the domain, responding to chemical cues and interactions with neighbouring cells. Initially, FBS is uniformly distributed, but as cells consume it at a constant rate, the concentration declines. At high FBS levels, cells aggregate. As FBS is depleted and its concentration decreases, aggregates break apart, mirroring experimental findings. Parameters related to cell size, motility, cell proliferation, and FBS consumption were inferred from experiments. Since calcium ions, key FBS components required for *Capsaspora* aggregation, are known to stabilise cell-cell adhesion (Ros-Rocher et al., 2023), we modelled FBS-dependent aggregation as an increase in adhesion strength. Specifically, our framework includes short-range repulsion to prevent cell overlap and longer-range FBS-dependent adhesion. Cells can interact and form adhesion sites via filopodia at a certain distance from each other, with adhesion becoming more pronounced in the presence of FBS. This assumption is supported by experimental observations showing FBS components within the cell body and along filopodia (Ros-Rocher et al., 2023).

Our model successfully reproduces all stages of FBS-induced aggregation. Within minutes of exposure to 5% FBS, cells form small clusters of 2-10 cells. Over the next 20-25 hours, aggregates grow through cell proliferation and merging during random migration. However, as aggregates enlarge, their movement becomes restricted due to stronger adhesion forces. Larger aggregates (hundreds of cells) primarily grow via proliferation rather than migration and only fuse when in close proximity. A key consequence of this behaviour is localised FBS depletion, which accelerates as aggregates grow (Figure 2b, bottom-right panels and Extended Data Figure 2b). The larger the aggregate, the faster it consumes FBS. Once depletion reaches a critical threshold, cells begin to disaggregate around 50-60 hours in experiments with 5% initial FBS. By 70 hours, cells are fully dispersed, indicating global FBS exhaustion. These processes are visualised in Movies 2-3.

Beyond capturing the experimentally observed aggregation-disaggregation dynamics, the model makes specific and testable predictions: If disaggregation is driven by FBS depletion then reintroducing FBS after complete disaggregation should rapidly restore aggregation, independently of the prior aggregation history, showing it is a reversible process dependent on FBS. Importantly, the model predicts that this response should occur within minutes (Movie 4), reflecting the immediate increase in effective cell-cell adhesion rather than slow genetic reprogramming. To test this prediction, we performed time-lapse experiments in which aggregation is induced with the addition of 5% FBS and subsequently allowed to disaggregate due to FBS depletion, after which 5%FBS is reintroduced into the medium at two different times, 24 h after disaggregation (Movie 4), and immediately after disaggregation (Movie 5). This behaviour was reproducible across independent replicates and persisted even after prolonged periods without FBS prior to serum re-addition, ruling out residual serum effects, and corroborating the full reversibility of the process predicted by the model.

Although individual realisations of our model vary slightly due to its stochasticity, and we do not account for medium flux effects on single-cell trajectories (see Supplementary Information), our model effectively captures *Capsaspora* aggregation dynamics. These results support the hypothesis that FBS stabilises cell-cell adhesion, driving multicellular aggregate formation. A key advantage of our model is its ability to visualise the FBS field in real time and explore different functional forms of FBS-induced adhesion, refining predictions beyond the assumption that FBS enhances adhesion strength.

### Threshold-like response to FBS

To better understand how FBS influences cell adhesion, we performed numerical simulations across a range of initial FBS concentrations. Experimental findings from a previous study (Ros-Rocher et al., 2023) indicate that at concentrations below 1% FBS, *Capsaspora* cells fail to form multicellular clusters, instead exhibiting random migration without a discernible pattern. However, when the FBS concentration reaches 1% or higher, aggregate size increases sharply, as reflected in the mean aggregate size dynamics previously reported (Ros-Rocher et al., 2023). This sharp transition suggests that FBS-induced aggregation in *Capsaspora* follows a threshold-like behaviour: adhesion remains minimal below 1% FBS, but once this threshold is exceeded, adhesion strength rises rapidly before reaching saturation at higher concentrations.

To capture this behaviour, we modelled cell-cell adhesion strength as a Hill function of the FBS concentration, whose sigmoidal shape reflects three key features: weak response at low FBS levels, a sharp increase in adhesion strength once the threshold is surpassed, and an eventual plateau at higher concentrations (Figure 3d). In addition to the Hill function, we tested two alternative adhesion models with linear responses to FBS. The first linear model assumed a gradual, moderate increase in adhesion strength with FBS concentration, while the second featured a steeper slope, producing a more abrupt adhesion increase (Figure 3d).

**Figure 3.**
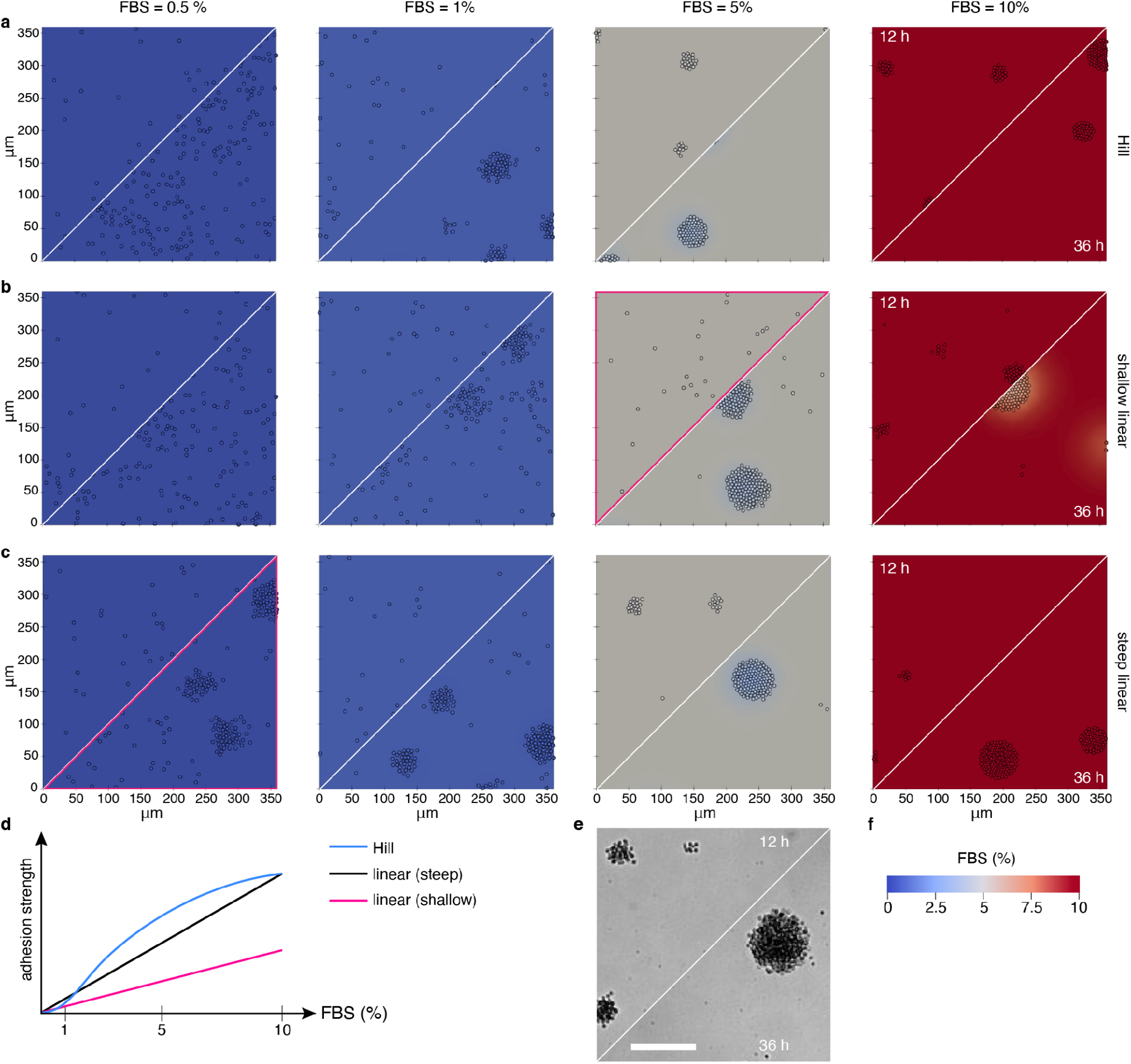
Comparison of different cell adhesion responses under varying initial FBS concentrations. We tested three functional form**s** for FBS-dependent adhesion: **a**, a sigmoidal Hill response, **b**, a shallow linear increase in adhesion strength, and **c**, a steep linear increase in adhesion strength whose functional forms are sketched in **d**. The initial FBS concentration, uniformly distributed at t=0 h, is indicated above each column, ranging from 0.5% (left) to 10% (right). To facilitate comparison, panel **e** presents experimental images from cultures with an initial 5% FBS concentration (scale bar: 100 μm). **f**, Figure legend for panels **a**-**c**. Each plot is divided diagonally by a white line: the upper half shows a snapshot at 12 h, while the lower half displays the system at 36 h. The colour of cells and the surrounding medium in simulations represents local FBS concentration, with red indicating high levels, grey intermediate levels, and blue low levels. Panels outlined in magenta highlight conditions that deviate from experimental observations and thus fail to reproduce the reported aggregation response to FBS in *Capsaspora*. Each image represents a single simulation, but we conducted five numerical simulations per condition, all yielding consistent results. The domain size for these simulations is 360 x 360 μm, with parameter values detailed in Extended Data Table 1. Full animations of plots shown in panels **a-c** are available in Movies 6-8, respectively.

Interestingly, and as shown in Figure 3a, the threshold-like adhesion response closely replicates experimentally observed *Capsaspora* aggregation dynamics. Specifically, at 0.5% FBS, no aggregates form; at 1%, small clusters emerge gradually over 36 h; and at 5% and 10%, aggregation occurs rapidly, consistent with experimental results. In contrast, the moderate linear response fails to capture the observed aggregation speed at 5% FBS (compare Figures 3b and e). In the model, adhesion remains too weak at 12 h to induce aggregation, whereas experimental observations show aggregate formation within minutes. The steep linear response reproduces aggregation at 5% and 10% FBS, but it also predicts aggregation at 0.5%, which was not previously observed (Ros-Rocher et al., 2023). Overall, these results support the hypothesis that *Capsaspora* adhesion to FBS follows a threshold-like response, where adhesion remains low below a critical concentration but increases sharply once this threshold is exceeded. To further validate this response, we quantified key metrics (area, cell number, and density) of the largest aggregate throughout our simulations. These measurements reinforce the conclusion that *Capsaspora* adhesion to FBS follows a threshold-like pattern (see Extended Data Figure 3).

Beyond influencing aggregation, the nonlinear upregulation of adhesion in response to FBS is also crucial to reproduce the aggregate disassembly dynamics. Notably, disassembly in large aggregates begins at the core of the aggregate, leading to an inside-out fragmentation pattern. This process occurs because aggregates develop with higher cell density at the centre, where adhesion and crowding effects are strongest. Consequently, FBS depletion occurs more rapidly in the core due to higher local cell consumption, while outer regions retain FBS longer, delaying disassembly. To visualise this process, we quantified local cell density in our simulations, colouring cells accordingly throughout the numerical experiment, and the results show disassembling at around 55 h, with the denser outer annulus persisting until approximately 70 h post-FBS induction (Movie 9).

### Temporal gene expression profiles reflect the dynamism of the aggregation process in Capsaspora

To explore the expression of the *Capsaspora* genome during aggregation, we first performed a clustering analysis of the expression profiles at different time points (Extended Data Figure 4). Principal component analysis (PCA) distinguishes two clear groups along the X axis, the first one formed by T0 (immediately before aggregation) & T1(3 min) clustering together representing adherent samples, and a second group including all the different stages of the aggregate formation after aggregation induction. The Y axis shows the clustering in order from T2 (12 h) to T7 (72 h) representing the aggregate growth. Based on sample distances, we differentiated four clusters, the first one formed by all the replicates from T2, T3 (24 h) & T4 (36 h) which cluster together representing the aggregative stage of *Capsaspora.* T0 represents the cell before inducing aggregation, also called “adherent stage”. T1 forms a cluster on its own which might be indicative of a specific response to the induction of aggregation. T5 (48h), T6 (60h) & T7 cluster as another group in between aggregative and adherent cells, which represents the disaggregation step in which we can find aggregates together with single cells in the culture. These results reveal that for 5% FBS, T5 (48h) should be considered part of the disaggregation step, situating the disaggregation process even earlier than what we previously anticipated by microscopy observations (55-60h from time lapses), and reveal that *Capsaspora* cells have an underlying regulatory mechanism to detect FBS depletion before the effects are observable in the culture.

Additionally, the DESeq2 analysis reveals that out of the 8352 genes analysed 922 genes are significantly upregulated at some point of the aggregation process and 990 genes are significantly downregulated, corresponding to approximately 11% and 12% of the analysed genes, respectively. Interestingly, T1 (collection beginning 3 min after aggregation induction) shows the highest number of up- and downregulated genes (5% in both cases), congruent with the fast response of *Capsaspora* to the addition of FBS.

To place these temporal expression changes in the context of the previously described *Capsaspora* life-stage transcriptomes, we reanalysed the RNA-seq data of Sebé-Pedrós et al. (2013) using the same current genome annotation and compared the transcriptional changes in the two studies. Because the two experiments were performed independently and no common biological sample was available to separate study-specific effects from biological differences, we primarily compared expression changes within each experiment rather than absolute expression levels between studies (Fig. 4a). For this comparison, we separately refitted both datasets using a common filtering and differential-expression procedure; the common inferential universe contains 8,002 genes. The original clustering and consecutive-time GO analyses were not recomputed with these new calls.

**Figure 4.**
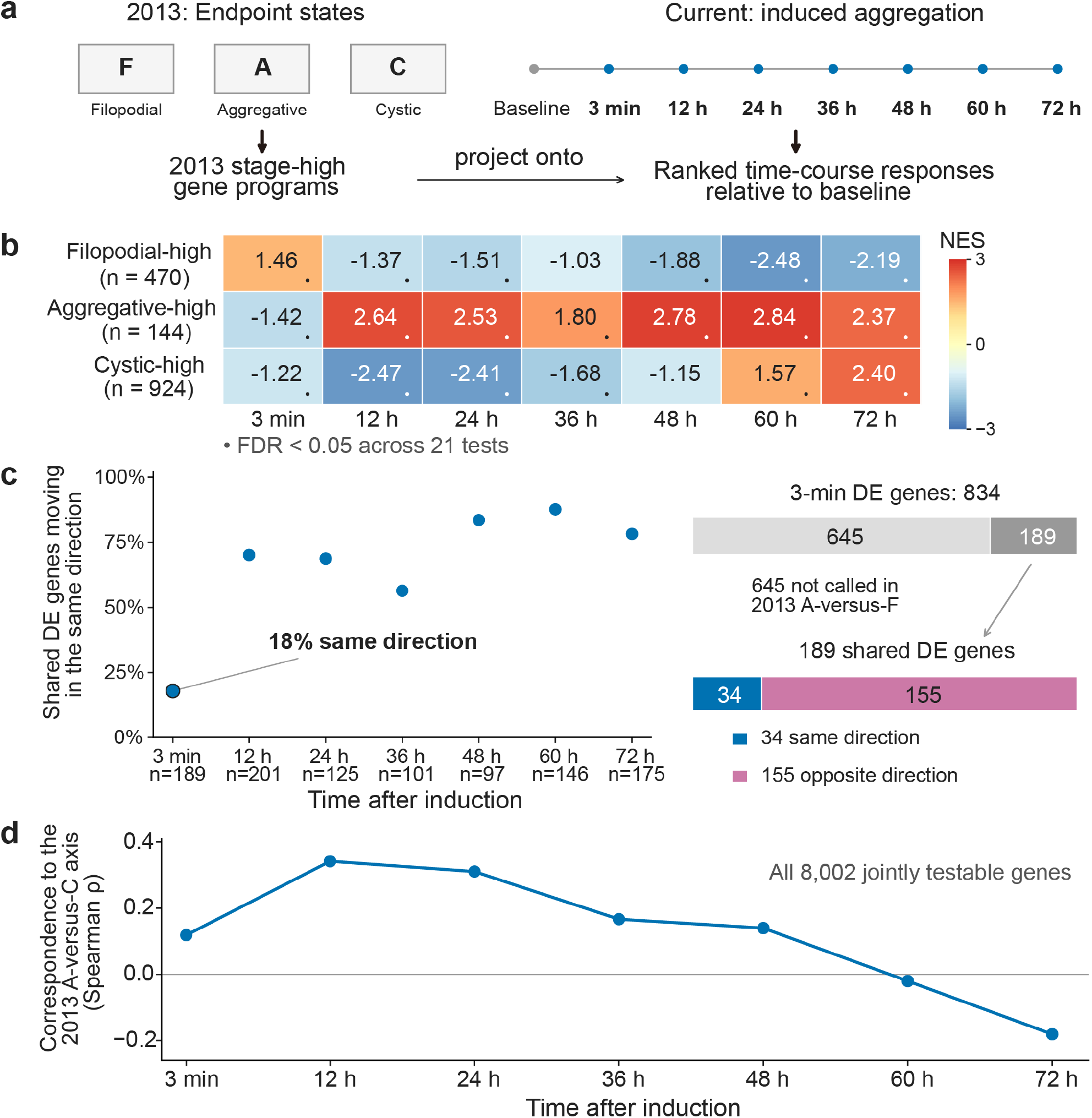
Temporal correspondence to reanalysed 2013 life-stage programs. (a) Study designs. F, A and C denote established filopodial, aggregative and cystic cultures. Comparisons use within-study changes because study and culture effects are confounded. (b) Preranked enrichment of stage-high gene sets, defined by significantly higher expression in one 2013 stage than in both other stages. Projected set sizes are F=470, A=144 and C=924 within 8,002 jointly testable genes. The normalized enrichment score (NES) indicates enrichment toward positive or negative time-course Wald statistics relative to T0; it is not a cell fraction or expression fold change. Dots indicate BH-adjusted FDR < 0.05 across 21 signature-by-time tests. Aggregative-high enrichment persists into the late phase alongside cystic-high enrichment. (c) Direction concordance among genes DE both at each current time versus T0 and in the 2013 A−F contrast. At nominal 3 min, 34 of 189 overlapping genes change in the same direction (18.0%) and 155 in the opposite direction. (d) Descriptive Spearman correlations of unshrunk log2 fold changes across the 8,002 genes. A−C correspondence is highest at 12-24 h, whereas C−F correspondence increases later. This differs from the timing of maximum aggregative-high NES in panel b. Time labels denote collection onset. T1 collection began at 3 min and was followed by 5-min centrifugation and handling before TRIzol lysis (Methods).

We first defined sets of genes that were expressed more highly in each of the three previously characterized states—filopodial, aggregative, or cystic—than in both of the other states. These stage-associated gene programs showed distinct behaviour during induced aggregation (Fig. 4b). At 3 min, genes characteristic of the filopodial state were preferentially increased, whereas aggregative- and cystic-associated genes tended to decrease. By 12-24 h, the aggregative-associated program was strongly enriched, coinciding with the formation and growth of multicellular aggregates. At later time points, particularly 60-72 h, genes associated with the cystic state became increasingly enriched. Notably, the aggregative-associated program remained detectable during this late phase, suggesting that disaggregation is accompanied by the emergence of a cystic-associated transcriptional program rather than a simple replacement of one discrete transcriptional state by another.

The nominal 3-min response was particularly distinct from the transcriptional changes previously associated with established aggregation. Among genes differentially expressed both at 3 min in our experiment and between the aggregative and filopodial states in the 2013 dataset, only 18% changed in the same direction (Fig. 4c). Moreover, most genes differentially expressed at 3 min were not called as differentially expressed in the previous aggregative-versus-filopodial comparison. Thus, temporal sampling reveals a prominent immediate response to aggregation induction that could not be resolved by an endpoint comparison of established life stages.

A complementary genome-wide analysis supported this temporal distinction (Fig. 4d). Expression changes separating the 2013 aggregative and cystic states showed their strongest correspondence to our 12-24 h samples and progressively weakened thereafter, becoming negatively correlated by 72 h. In contrast, transcriptional differences between the cystic and filopodial states became increasingly similar to the late phase of our time course. Together, these analyses indicate that the previously described aggregative transcriptome contains multiple temporally separable components that are resolved by following aggregation and disaggregation dynamically.

To identify temporal expression patterns in the original analysis, we used degPatterns from DEGreport (Pantano), followed by hierarchical grouping into six super groups (Figure 5; Extended Data Figure 5). The consensus-index CDF was used to guide this grouping. We then examined GO term enrichment within each group. This grouping summarizes expression profiles rather than defining discrete developmental stages.

**Figure 5.**
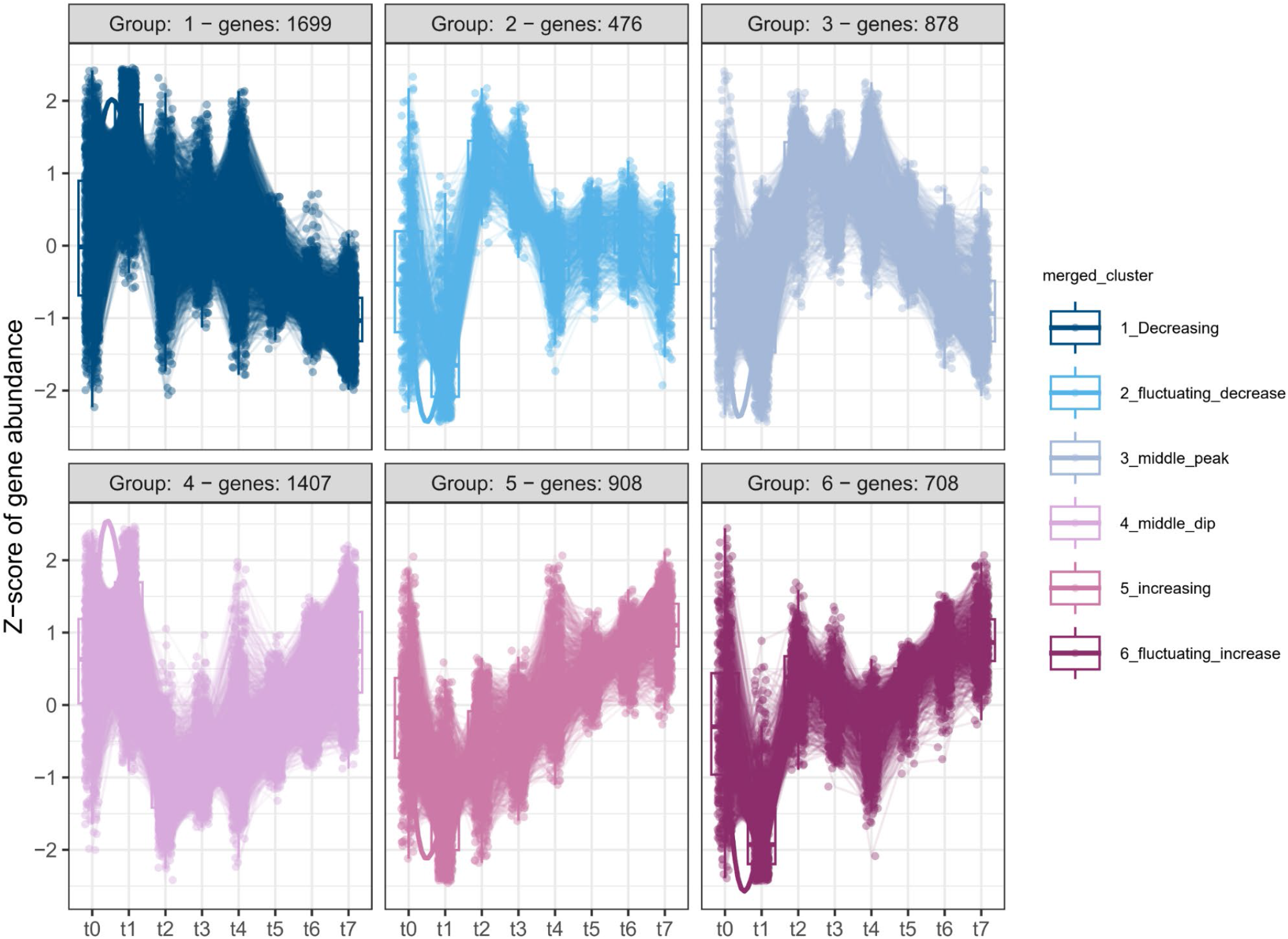
Gene expression dynamics are captured in six super groups. Decreasing pattern. (Group 1), in which genes keep decreasing their expression values after 12h. In this group we find genes related to cell growth and division, mainly the regulation of mitosis and the ribosome biogenesis. The **Fluctuating Decreasing** group (Group 2) shows genes whose expression levels decrease at 12h followed by an increase in expression and a new decrease after 24 h. This group includes GO terms related to the cell cycle process, the regulation of actin polymerization, as well as GO terms related to multicellular functions such as the positive regulation of Notch pathway, system development, and morphogenesis. The **middle-peak** group (Group 3) reflects GO terms that are especially relevant in the maintenance of the already formed aggregates, such as post-translational modifications and splicing. The **middle-dip** group (Group 4) includes genes whose expression levels are the lowest in the aggregates that are fully formed. Here, we find GO terms related to sterol metabolism, gluconeogenesis and ubiquitination processes. The **Increasing group** (Group 5) shows a smooth increase in the expression levels across all time points after 12h, and includes GO terms related to Development and signalling, cell fate determination, migration, proliferation, the regulation of stress (oxidative stress HIF1, and osmotic), and the negative regulation of cell population signalling. The **Fluctuating-increasing** group (Group 6) depicts an increasing trend after 12h of aggregation, and includes GO terms related to development & growth, the positive regulation of multicellular organ formation, cellular localization, actin cytoskeletal organization, and the negative regulation of the Hippo pathway.

To identify and extract gene expression clusters we used DEGpatterns (Pantano) and observed 52 groups of genes. The cumulative distribution function (CDF) of the consensus index indicates that 6 clusters of different expression patterns are the minimum to capture all the variation in gene expression (Extended Data Figure 5). Thus we grouped the 52 patterns into 6 super groups based on hierarchical clustering (Figure 5) and performed a GO term enrichment analysis to determine the main functions associated with each expression profile.

### Main functions underlying the aggregation process in Capsaspora

In addition to the analysis of overall temporal gene expression patterns mentioned above, we performed differential gene expression analysis between consecutive time points to identify genes responsible for the transitions between each stage. A striking feature is the fast response of *Capsaspora* to the addition of FBS and induction of aggregate formation. In the sample whose collection began at 3 min, we observed the highest number of genes being up- and downregulated, in which the cell rapidly favours cell growth and proliferation, with an increase in GO terms related to ribosome biogenesis and growth, while genes related to transcription are downregulated. This first response is followed by a period of formation of small aggregates during the first 12 h in which we observed an increase in GOs related to development and regulation of cell cycle and cell-cell signalling, with a reduction of genes related to ribosome and amino acid metabolism (Figure 6 and Extended Data Figure 6). During the period that corresponds to the 12 to 24 h in which aggregates have reached larger sizes, we find genes related to lipid, glucose and amino acid metabolism, which suggest functions related to growth and metabolism within aggregates. Between 24 and 36 hours the aggregates keep merging with each other whenever they come into contact with one another. In this period, we also observed downregulation of genes related to actin-based processes and cytoskeletal organization, which could be related to the depletion of FBS in the media (as also predicted by our model simulations; see Movies 2-3), and the initial steps of the disaggregation signal. In the period between 48-60 h, we find upregulation of GO terms related to Golgi vesicle transport and the downregulation of GO terms related to ribosome biogenesis (which is highly energy demanding for the cell), amino acid synthesis and growth, probably indicative of cells being under stress within the aggregates, or that they could even be entering into cyst formation. The final period between 60 and 72 h corresponds to the complete dissociation of the aggregates in which we find the downregulation of GO terms related to the nucleosome organization.

**Figure 6.**
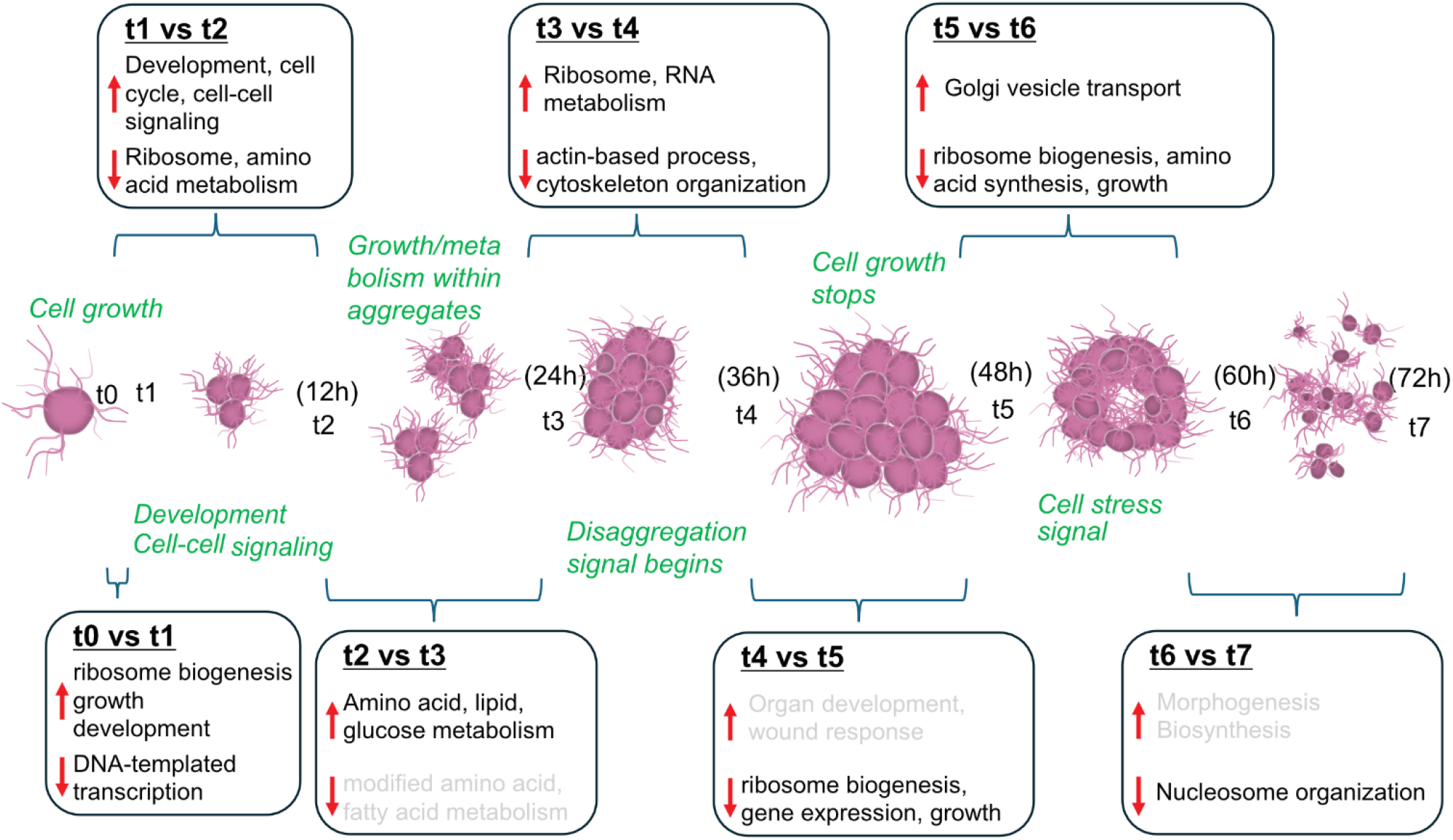
GO term enrichment analysis of DE genes during *Capsaspora* aggregation. The aggregation process was divided into 7 key moments that reflect its dynamism. GO enrichment analyses of DE genes were performed between consecutive time points, black GO terms are significantly enriched (p<0.05) while grey GOs present higher p values.

In conclusion, as expected, the first two steps of the aggregation process are those with the largest deployment of genes, revealing the importance of the genetic mechanisms regulating the first response to FBS and the initial formation of the aggregates. Once the aggregates are formed, most of the differences are related to metabolism and cell cycle. The other peak of DE genes is at T4 vs T3 when genes related to disaggregation start to be expressed which correlates with FBS depletion.

Additionally, to discern specifically which genes are being deployed, we looked at the expression dynamics of genes that are particularly relevant for animal multicellularity, including cell adhesion molecules such as integrins, cadherins and the Dystrophin-associated glycoprotein complex (DGC), development-related transcription factors, receptor TKs and organ growth control components (Extended Data Figure 7).

In animals, ECM-cell adhesion is regulated by the interaction of integrins and associated cytoplasmic proteins such as vinculin. Our data reveal the upregulation of two out of the four integrin B present in *Capsaspora,* specifically both integrin B1 together with vinculin have a peak in expression at 24 h of aggregation after which they decrease recovering their basal levels after disaggregation. This dynamic fits with our model predictions in which FBS concentration would regulate the aggregation dynamics and disaggregation signalling starts earlier than observed. Another interesting integrin is integrin B2, which is expressed in *Capsaspora* filopodia (Parra-Acero et al., 2020), and that presents a fast, strong response to the addition of FBS and its expression levels keep increasing during the entire process of aggregation. These results are also consistent with the expression pattern of many *Capsaspora’s* cytosolic tyrosine kinases homologous to the animal counterparts (Suga et al., 2012), which peak in their expression values at 12 h, once the aggregates are formed. Other metazoan relevant ECM-cell adhesion mechanisms like the Dystrophin-associated glycoprotein complex (DGC) display a very fluctuating pattern of expression. With regard to cell-cell adhesion, Cadherins and C-type lectins are downregulated during the formation and growth of the aggregates and recover their regular levels of expression after disaggregation.

Regarding development-related TFs, bZIP and TBox transcription factors (TFs) are among the most fluctuating TFs found in *Capsaspora.* Other interesting TFs are Runx 1 & 2 both of which are rapidly expressed upon the addition of FBS. Mef2 and p53 show a fast downregulation during the first moments of aggregation. NfKB is downregulated during the formation and maintenance of the aggregates, recovering its expression levels during disaggregation, which might be indicative of their implication in cell proliferation or stress response programs.

Another key developmental program in metazoans is the Hippo pathway. We find most of its core regulatory components to be dynamically expressed during aggregation, with the peak expression at 12 h or 24 h. For example, Merlin and Kibra are upstream regulators of the pathway involved in mechanical contact sensing. Both are highly upregulated during aggregate growth and downregulated during the disaggregation phase. One of the main components, Warts/Lats, shows a different pattern where it remains unchanged or low during aggregation. But we do find the expected upregulation during aggregate growth for most other components including Hippo/Mst and Yorkie/Yap. Interestingly, most components show upregulation during the disaggregation phase, after being downregulated at around 36 h.

We also found *Capsaspora* homologs of animal heterotrimeric G-protein α subunits, “Gαv” & “Gαi/t/o”, that display a positive fast response upon the addition of FBS to the media, which could be indicative of the detection of FBS and trigger of aggregation. In addition to the G-protein subunits homologous to those of animals, we found potential G receptors specific to *Capsaspora*, (COWC_0686, COWC_090031, COWC_01017) that share the same response dynamic making them putative targets for FBS detection and the initial trigger of aggregation.

Overall, these results indicate that *Capsaspora*, a close unicellular relative of animals, already displays a dynamic regulatory mechanism involving genes that are relevant to animal multicellularity, which is an indication that cell aggregation could have had a role in the origin of those “multicellular genes” that later on were co-opted to work within a multicellular system.

## Discussion

Building on the *Capsaspora* life-stage transcriptomes described in 2013 (Sebé-Pedrós et al., 2013), this study resolves expression changes across serum-induced aggregation and disaggregation and supplies a complementary quantitative model of the morphological dynamics. Its principal advance is temporal resolution, including an immediate response distinct from the established aggregative-versus-filopodial comparison.

The sample whose collection began at 3 min shows an extensive change in RNA abundance. Collection included a further 5-min centrifugation before TRIzol lysis, so these differences cannot be assigned to the first 3 min after induction. RNA-seq measures relative RNA abundance rather than nascent transcription, so rapid transcription, RNA stability and sample handling cannot be distinguished here. The comparison with 2013 data is consistent with a serum-response component but cannot establish that serum or nutrients dominate the response. Recent work shows that initial aggregation can occur despite inhibition of new protein synthesis and implicates lipid endocytosis and filopodia retraction (Kidner et al., 2026). This supports separating the physical initiation of aggregation from subsequent transcriptional dynamics; it does not determine the origin of the RNA changes in our experiment.

Comparison with the previously described *Capsaspora* life-stage transcriptomes further clarifies what is gained by temporal sampling. The transcriptome of established aggregative cultures described by Sebé-Pedrós et al. (2013) does not correspond to a single point in our induced time course. Instead, its different transcriptional components show distinct temporal behaviour. The genome-wide pattern distinguishing aggregative from cystic cells is most evident at 12-24 h, whereas cystic-associated expression becomes increasingly apparent during the late disaggregation phase. Most strikingly, the response in the sample collected starting 3 min after FBS addition is poorly represented by the transcriptional changes observed in established aggregates, revealing an immediate induction program that could not have been detected by the previous endpoint experimental design. Thus, temporal resolution refines the earlier description of three *Capsaspora* life stages by revealing the transitions and partially overlapping regulatory programs connecting them.

Our modelling study suggests that FBS-upregulated cell adhesion is a possible mechanism underlying *Capsaspora* aggregation dynamics. By calibrating our model with experimental data, we successfully captured all stages of the process, from initial clustering to aggregate growth and eventual disassembly following FBS depletion. Moreover, our results indicate that *Capsaspora* adhesion exhibits a nonlinear, threshold-like response to FBS: minimal adhesion at low concentrations, a sharp increase past a critical threshold, and saturation at higher levels, allowing the prediction of the reversibility and reproducibility of the process.

The success of our model using a Hill function with a coefficient of approximately 2 may also be considered a prediction about the underlying biology. A coefficient of this magnitude is a classic signature of cooperative processes, such as receptor dimerization for stable adhesion. This is intriguingly consistent with our transcriptomic data showing the dynamic upregulation of integrins, which function as obligate heterodimers. It is noteworthy, moreover, that only when a non-linear FBS-dependent cell adhesion function is considered does our model reproduce the experimental results and observations for cluster size distributions, the disaggregation dynamics, or the threshold-like behaviour of the onset of cluster assembly. A linear response function fails to reproduce the details of the (dis)aggregation process. A relevant question is whether this cooperative behaviour arises from synergistic interactions between the multiple components in FBS or from the intrinsic properties of the cell’s own sensory and adhesion machinery. Recent evidence supports the latter possibility. A recent study has shown that *Capsaspora* exhibits a similar sharp, threshold-like aggregation response when induced by a single, pure synthetic lipid, dioleoyl phosphatidylcholine (DOPC) (Kidner et al., 2024). This finding suggests that the cooperative nature of the response is an inherent feature of the cellular system itself, likely involving mechanisms such as receptor clustering. The precise molecular nature of these receptors remains unknown and may be an important direction for future work.

Mathematical modelling has been previously used to describe aggregation of cells (or larger species) in various contexts. Our model aligns most closely with non-local adhesion models (Buttenschön & Hillen, 2021) (Armstrong et al., 2006) (Griffié et al., 2020) (Noureen et al., 2025), which describe aggregation driven by cell adhesion. While these models are often used to explain cell sorting through differential adhesion strengths, they have also been applied to multicellular cluster formation (Armstrong et al., 2006) (Griffié et al., 2020; Noureen et al., 2025). Similarly to these models, our approach incorporates random motility and cell-cell adhesion but is specifically calibrated for *Capsaspora*. In contrast, other commonly used clustering models do not align with *Capsaspora* behaviour. In particular, *Capsaspora* cells do not exhibit coordinated or persistent movement (see Extended Data Figure 9 and Supplementary Information), ruling out swarming-type models such as Vicsek and Cucker-Smale (Cucker & Smale, 2007; Vicsek et al., 1995) and self-propulsion-based models that generate multicellular clusters through motility-induced phase separation (Caprini et al., 2020; Gonnella et al., 2015). Additionally, chemotaxis-based models that can reproduce clustering (Avery et al., 2021; Keller & Segel, 1970; Palsson & Othmer, 2000) are also inconsistent with *Capsaspora* aggregation, as there is no evidence of chemical signalling between cells, nor do they move up an FBS gradient during (dis)aggregation. Chemotactic behaviour during (dis)aggregation is not observed in the experiments, as can be seen in Supplementary Movies 1 and 2. This observation is further supported by the fact that our mathematical model, which does not account for chemotaxis, can reproduce the whole dynamics of aggregation and disaggregation, including cluster size quantification, timing and dynamics of cluster disassembly, and threshold behaviour with respect to FBS concentration. Our model provides a detailed, agent-based framework that captures the behaviour of individual *Capsaspora* cells with parameters calibrated from experimental data. Crucially, our model explicitly couples cell-cell interactions with the temporal dynamics of FBS, allowing aggregation to emerge as a dynamic and reversible process controlled by extracellular FBS availability. This coupling explains why larger aggregates dissociate from the centre while smaller aggregates merge more rapidly under higher FBS concentrations, and is essential for reproducing the reversible nature of *Capsaspora* aggregation. Directly measuring FBS concentration at different stages of aggregation would allow a more detailed mapping between model variables and experimental observables.

Nevertheless, these predictions are indirectly supported by the GO enrichment analysis of differentially expressed genes during the aggregation process in which GOs related to growth, development, and cell-cell signalling among others are highly enriched in the first time points after the aggregate induction, reflecting the high diffusion rate of FBS in the media and the fast response in *Capsaspora.* Furthermore, a significant decrease in GO terms related to actin-based processes, cytoskeletal organization and growth is captured between 24 and 36 h of aggregation, suggesting that the onset of the disaggregation starts earlier than observed in the microscopy imaging. Finally, consistent with model predictions, aggregation remains fully reversible: after dissociation at later stages, re-addition of FBS triggers a rapid reaggregation response within minutes as anticipated from the model and in agreement with previous experimental observations (Ros-Rocher et al., 2023).

This chemically regulated aggregation-disaggregation scenario also couples with previously established hypotheses in which morphological variation in response to the environment is an ancient physical property already present in the last unicellular common ancestor. This may have been especially relevant in earlier multicellular forms where the underlying regulatory networks were not yet fully established, and the interplay of intrinsic physical properties and external conditions was likely more prominent. Such interplay is also found in other phylogenetically unrelated aggregative lineages such as dictyostelids, fungi or even myxobacteria (Arias Del Angel et al., 2020; Newman et al., 2006), and may have provided a template for the accumulation of stabilizing and reinforcing genetic circuits that ultimately led to the development of obligate animal multicellularity.

The rapid and dynamic nature of *Capsaspora* aggregation might be indicative of or reflect their biological importance for unicellular organisms, which need to adapt very fast to changing environments, and could have played a crucial role in the early evolution of multicellularity by conferring an adaptive advantage in fluctuating environments. *Capsaspora* has only been found inside the freshwater snail *Biomphalaria glabrata* (Stibbs et al., 1979) which could suggest that the aggregation response is a mechanism to protect against the host response, but *Biomphalaria* does not have the classical vertebrate lipoproteins used in FBS, which could imply that *Capsaspora’s* response is an evolved response to chemical cues from the snail that can indicate the identity and physiological state of its host (Kidner et al., 2024). Alternatively, *Capsaspora* could also live outside the snail and the signal would come from other neighbouring organisms. This could be of biological relevance since passive aggregation or autoaggregation, can be influenced by the density of surrounding single cells. When competition between aggregates and single cells is low, aggregates experience a growth disadvantage due to restricted access to resources in their interior. However, under high competition, aggregates exhibit increased fitness, as vertical extension above the surface provides cells at the top with improved access to nutrients (Trunk et al., 2018). Additionally, aggregation may enhance *Capsaspora*’s dispersal capability, facilitating colonization of new locations (Kragh et al., 2016; Tong et al., 2022).

Our transcriptome analyses reveal the dynamic expression of relevant genes such as integrin and several tyrosine kinases and TFs associated with multicellular functions in metazoans during the aggregation process, as it happens with *Ministeria vibran*s (Li et al., 2026). The fact that those genes are being dynamically deployed during aggregation and that some of them originated at the Filozoa clade (Metazoa, Choanoflagellata, and Filasterea), suggests that those genes could have originated to function for these facultative aggregation processes, providing an important raw genetic material that animals could later co-opt to work within obligate multicellular entities. If so, this situates aggregation as a relevant mechanism involved, directly or indirectly in the origin of animals. This is in contrast to the classical views that cell aggregation never gives rise to complex multicellularity. It should be mentioned that those are *ad hoc explanations*, since they are based on the observation that extant complex multicellular taxa (i.e., animals, plants, fungi, red and brown algae) develop by clonal division, while some (not all) extant simple multicellular taxa (i.e. *Dictyostelium, Acrasis*) develop through cell aggregation. However, whatever the development of extant organisms nowadays (clonal or aggregative) cannot be taken as a proxy for how the unicellular-to-multicellular transitions initially took place.

Our data suggest that chemically induced cell aggregation, highly influenced by cell adhesion, was most likely present in the unicellular ancestor of animals and suggests an ancient, environmentally responsive property that may have served as a template for more complex, genetically stabilized multicellularity. During cell aggregation, the unicellular ancestor may have used many genes that later became pivotal for animal multicellularity and development. Thus, the hypothesis that the first animal emerged through cell aggregation and subsequently evolved into an obligate multicellular entity with embryonic development cannot be ruled out (Ruiz-Trillo et al., 2023).

Our work, notably the development of a mathematical model that consistently recapitulates all the features of the aggregation process, firmly establishes *Capsaspora* as an integrative and tractable model system to further analyse the evolutionary potential of cell aggregation to either evolve genes that later became crucial to animals or even to create the first animal. In particular, having this mathematical model, together with the recent establishment of transfection and future optimization of genome editing tools in *Capsaspora*, will allow researchers to interrogate the potential ancestral functions of many genes central to animal multicellularity by making it possible to quantitatively assess subtle phenotypic consequences that would otherwise remain hidden.

## Materials and Methods

### Cell culture and aggregate induction

*Capsaspora owczarzaki* cells (strain ATCC 30864) were maintained axenically in ATCC medium 1034 (modified PYNFH medium) with the addition of 10% (v/v) fetal bovine serum (Sigma-Aldrich, F9665) at 23°C.

Aggregate formation was induced chemically following a protocol adapted from (Ros-Rocher et al., 2023). Cells were washed twice in FBS-free growth medium by centrifugation at 5000g for 5 min. Subsequently, 1×10^6^ *Capsaspora* cells were seeded in a 12 well plate (Costar® 12-well plates REF3513) coated with collagen to generate a low-adhesion surface for *Capsaspora* (Parra-Acero et al., 2020)

Collagen coating was performed by adding 600µl of 20µg/mL collagen into each well for a minimum of 30 min, after which it was removed and the wells were left to dry completely. Cells were then seeded in FBS-free medium and incubated overnight. Aggregation was then induced by adding FBS at the desired concentrations.

Bright-field imaging was performed using a Zeiss Axio Observer Z.1 epifluorescence inverted microscope equipped with LED illumination and an Axiocam 503 mono camera at 5x magnification for (Supp Videos 1,2/ 5&10%FBS aggregation), or an Eclipse TS100 Nikon epifluorescence inverted microscope equipped with an Intensilight C-HGFI Illuminator and a DS-Fi2 Camera Head at 10× magnification (Supp Video3/0,05%FBS aggregation). Average aggregate areas were measured by batch processing with a macro script in Fiji imaging software version 2.14.0/1.54f

### Computational model of Capsaspora aggregation

We developed a computational model to investigate the mechanisms of *Capsaspora* aggregation. Our model consists of two key components: *(i)* an agent-based cell model to account for *Capsaspora* proliferation and migration (Extended Data Figure 1a) and *(ii)* a continuum description of the dynamics of Fetal Bovine Serum (FBS), a chemical stimulant in the culture medium which can affect *Capsaspora* behaviour (Extended Data Figure 1b). These components are coupled to simulate cell behaviour under varying FBS concentrations (Extended Data Figure 1c). The model has been calibrated using experimental data where available, ensuring that key processes, such as cell proliferation, FBS-dependent motility, and FBS dynamics, are informed by observed *Capsaspora* behaviour. Here, we summarise the main processes incorporated in our *Capsaspora* model, while Supplementary Information provides a detailed description of the model.

#### Cell model

The centre-based approach employed in this work treats each cell as a circular entity in two dimensions (Extended Data Figure 1a). Adult (fully grown after cell division) cells are assumed to have fixed cell radius and thus, we only need to track the position of their centres to know the distribution of cells across the domain. The position of the centre of *i* th cell, *x_i_*, evolves according to the following ordinary differential equation:

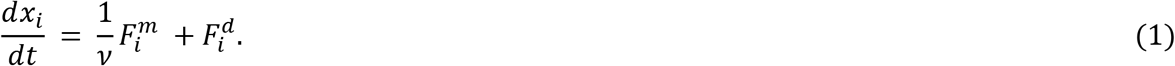

In this equation, *ν* represents the drag coefficient, and 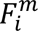 denotes the mechanical forces arising from interactions with neighbouring cells. These forces include short-range repulsion when cell centres are in close proximity and long-range adhesion mediated by *Capsaspora* filopodia (Extended Data Figure 1a). As shown in (Ros-Rocher et al., 2023), the metabolic activity of Capsaspora in the presence of externally supplied FBS increases the ‘stickiness’ of their filopodia, enhancing long-range interactions between cells. Accordingly, we assume that the strength of these adhesion forces increases with local FBS concentration.

In addition to mechanical forces, cell movement is influenced by random motility, represented by the term 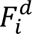 in Eq (1). We calibrated the random motility component using experimental data on individual cell trajectories at various FBS concentrations. Our analysis, detailed in Supplementary Information, demonstrates that *Capsaspora* motility is significantly enhanced at higher FBS levels. To capture this behaviour, we model cell motility as an increasing function of FBS concentration.

Thus, *Capsaspora* aggregation observed in experiments can arise in our model simulations due to a balance between FBS-mediated adhesion and random motility. In the absence of FBS, *Capsaspora* cells fail to form aggregates (Ros-Rocher et al., 2023), which suggests that the adhesion forces are weaker than the random motility that disperses the cells. However, when stimulated with FBS, aggregation occurs, indicating that the FBS-dependent cell interactions dominate, even though cell motility also increases in high-FBS environments.

In our simulations, we also incorporate cell proliferation, modelled stochastically based on a distribution fitted to experimental data on *Capsaspora* division times, as reported in (Pérez-Posada et al., 2020). Cell division is assumed to be symmetric, consistent with the findings in (Pérez-Posada et al., 2020), where each division produces two daughter cells of approximately equal size. After division, the daughter cells gradually grow to reach the size of an adult *Capsaspora* over a specified period. Cell death is not accounted for in the computational model, as, in the time scale of the experiments, cell death is negligible. We refer the reader to the Supplementary Materials, Sections I.1 and I.2 for a full account of the details of our model implementation.

#### FBS model

The FBS dynamics are modelled by a partial differential equation (PDE) (Extended Data Figure 1b), which describes its diffusion across the medium and consumption by the cells. The diffusion coefficient and uptake rates were fitted based on experimental data on FBS depletion, which leads to *Capsaspora* disaggregation behaviour.

#### Coupling between cell and FBS models

The model couples cell behaviour with FBS concentration (Extended Data Figure 1c), allowing motility and intercellular adhesion to vary with local FBS levels while cell metabolic activity depletes the available FBS. This coupling enables the simulation of cell aggregation patterns as a response to varying FBS concentrations, providing insight into the mechanisms driving *Capsaspora* aggregation.

### RNA extraction and Temporal transcriptomics analysis

Total RNA was extracted from the cells in triplicate for each condition after 0 min (i.e., immediately before aggregate induction, T0), 3 min (T1), 12 h (T2), 24 h (T3), 36 h (T4), 48 h (T5), 60 h (T6) & 72 h (T7) of incubation, using TRIzol (Invitrogen/Thermo Fisher Scientific, 15596026) as described in (Ocaña-Pallarès et al., 2019). RNA was purified with QIAGEN RNeasy Mini Kit (ref 74104). RNA concentration was measured using a Qubit 2.0 fluorometer from Invitrogen. Samples were sent to NOVOGENE for sequencing on Illumina NovaSeq 6000 platform with paired-end 150 bp sequencing strategy, library preparation was performed by polyA capture and cDNA synthesis, purification was checked by PCR. Time labels denote the start of cell collection after induction, rather than the time of RNA stabilization. Cells were collected by centrifugation at 5,000 g for 5 min; medium was removed and the pellet lysed in TRIzol. Thus, T1 collection began at 3 min, but RNA stabilization occurred after the additional centrifugation and handling interval. The precise induction-to-lysis interval was not recorded here.

Differential gene expression analysis was performed with DESeq2 (threshold 0.05) (Love et al., 2014). Identification and visualization of gene expression clusters was performed with DEGpatterns (Pantano), and out of the 52 clusters obtained we determined the best number of consensus clusters based on the cumulative distribution function (CDF) of the consensus index vs the number of clusters resulting in the 6 consensus clusters represented in Figure 5 (Pantano). The GO term enrichment analysis and annotation was performed with topGO (Alexa & Rahnenführer). A detailed script of each of the steps has been uploaded in our GitHub repository.

### Harmonised reanalysis for the 2013 comparison

For Figure 4 only, 18 public runs (SRR515464-SRR515481; PRJNA20341) were grouped into nine biological libraries, three each for filopodial, aggregative and cystic cultures. Two unreplicated 2010 runs were excluded. The 2013 reads were trimmed with fastp (Phred threshold 15; minimum length 40 bp) and quantified against the v5 CDS reference using RSEM with Bowtie 2. Gene-level expected counts were obtained by collapsing transcript suffixes. Libraries were quantified as unstranded for comparison with the available current-study count matrix; a strand-aware sensitivity check on one library per 2013 stage gave gene-level Spearman correlations of at least 0.993. The current time course was refitted from its existing count matrix, not newly sequenced or fully remapped.

Genes with *counts per million* (CPM) ≥ 1 in at least three libraries within each study were retained, giving 8,319 genes in 2013 and 8,206 in the current study. Expected counts were rounded after transcript aggregation for 2013 and before aggregation for the current matrix, matching the audited input processing. Separate DESeq2 models used stage or categorical sampling time, with filopodial or T0 reference levels. Replicate numbers were not treated as paired blocks. Two-sided effect-thresholded Wald tests used |log2 fold change| > 0.585 and IHW-adjusted P < 0.05 within each contrast. This reanalysis yielded 1,797 unique current-study DE genes across seven baseline contrasts (920 ever up, 972 ever down, 95 in both directions). It was not used to replace the original clustering or interval-specific GO analyses.

Cross-study inference used 8,002 genes passing expression filtering with finite test values in both studies. Of 837 DE genes at 3 min in the full current-study reanalysis, 834 belong to this shared universe. Stage-high sets required higher expression in one 2013 stage than in each of the other two under the same effect-thresholded criteria. Shared-set sizes were 470 filopodial-high, 144 aggregative-high and 924 cystic-high genes. These are reanalysis-defined signatures, not the original published lists.

For each current-study time versus T0, all 8,002 genes were ranked by the zero-null DESeq2 Wald statistic, with gene identifiers breaking ties. fgseaMultilevel used minSize=15, maxSize=5000, eps=0, nPermSimple=10000, sampleSize=101 and SerialParam, with seeds 20260922-20260928. Benjamini-Hochberg correction was applied across the 21 signature-by-time tests. Direction concordance used genes DE in both comparisons; genome-wide Spearman correlations used unshrunk within-study log2 fold changes and are descriptive. Models used R 4.5.3, DESeq2 1.50.2 and IHW 1.38.0; signature enrichment used fgsea 1.36.2. Categorical contrasts do not fit a continuous temporal trajectory. Independent study and culture effects remain inseparable from biological differences without shared reference samples.

### Use of generative artificial intelligence

Claude (Anthropic) and Codex (OpenAI) were used to assist with planning revisions and editing the manuscript. AI assistance also supported the development and checking of analysis scripts and the synthesis of results from the comparative reanalysis of the *Capsaspora* RNA-seq datasets, including the Sebé-Pedrós et al. (2013) dataset. The authors retain responsibility for the analysis methods, verification of code and outputs, interpretation of results, and all statements in the manuscript.

## Data availability statement

The simulation results presented in this paper have been uploaded to FigShare at https://doi.org/10.6084/m9.figshare.28761560. Using the scripts provided in our GitHub repository (https://github.com/daria-stepanova/Capsaspora.git), all figures and movies from this study can be fully reproduced.

The Movies 1 to 9 presented in this paper have been uploaded to FigShare at https://doi.org/10.6084/m9.figshare.29044817.v4.

The transcriptomic raw data used for the analyses during the current study are available in the EMBL-EBI repository with the accession code PRJEB89428.

The *Capsaspora* annotated genome used for the analyses is available at the DDBJ repository under the BioProject: PRJDB19057 (PSUB024242)

## Code availability statement

The code for our mathematical model is available on GitHub at https://github.com/daria-stepanova/Capsaspora.git. The repository includes a detailed README file with instructions on running the model and reproducing the simulation results presented in this study.

## Acknowledgements

Work in the IRT lab is supported by grants PID2020-120609GB-I00 funded by MICIU/ AEI /10.13039/501100011033/ and by “ERDF A way of making Europe”, and by PID2023-153273NB-I00 funded by MICIU/AEI /10.13039/501100011033 and FEDER, UE. We also acknowledge support to Departament de Recerca i Universitats de la Generalitat de Catalunya (exp. 2021 SGR 00751) and support by PIE-202120E047-Conexiones-Life. D.S. and T.A. thank the CERCA Program/Generalitat de Catalunya for institutional support. D.S. and T.A. have been funded by grant PID2021-127896OB-I00 funded by MCIN/AEI/10.13039/501100011033 ‘ERDF A way of making Europe’. The work of D.S. and T.A. has been supported by the Spanish Research Agency (AEI), through the Severo Ochoa and Maria de Maeztu Program for Centers and Units of Excellence in R&D (CEX2020-001084-M). K.K. received the support of a fellowship (LCF/BQ/PI20/11760009) from “la Caixa” Foundation (ID 100010434) and from the European Union’s Horizon 2020 research and innovation programme under the Marie Skłodowska-Curie grant agreement No 847648, as well as JSPS KAKENHI Gran-in-Aid for Research Activity Start-up (24K23213) and Grant-in-Aid for Transformative Research Areas (A) (26H01826).

## Author contributions

All authors contributed to the conception of the project. D.S. and T.A. developed the computational models. D.S. performed all the simulations. G.B.S. captured videos of *Capsaspora* aggregation. K.K. and G.B.S. performed the RNA-seq experiments. G.B.S. analysed all the RNA-seq data. G.B.S., D.S., and I.R.T. prepared the first draft of the manuscript. K.K., T.A. and I.R.T. were involved in supervision and funding acquisition. All authors contributed to the revision of the manuscript.

## Additional information

Supplementary information and videos accompanying the manuscript are available at FigShare (https://doi.org/10.6084/m9.figshare.28761560.v1).

## Extended Data Figure and Table legends

**Extended Data Figure 1.**
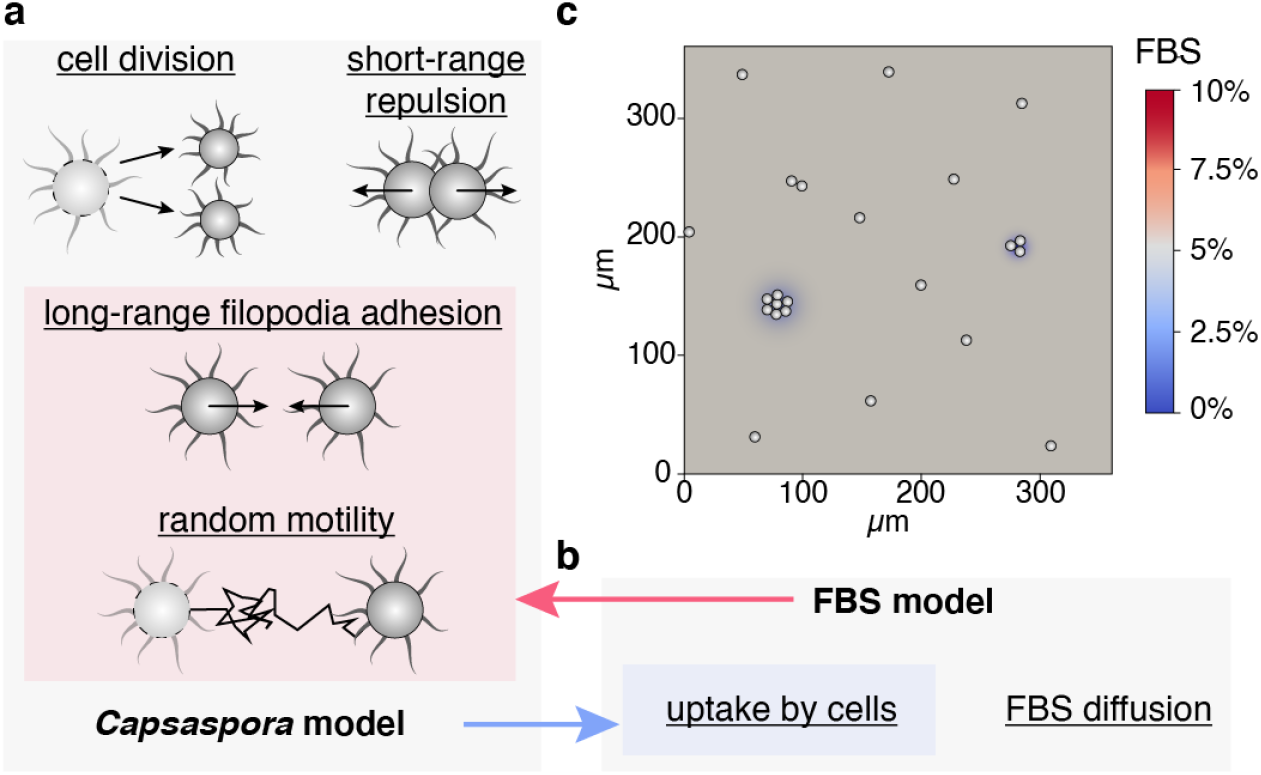
A schematic representation of our modelling approach. **a,** A centre-based model is used to describe the behaviour of proliferating and migrating *Capsaspora* cells. In two dimensions, each cell is modelled as a circle of radius, *r*, and the positions of cell centres are tracked throughout the simulations. Cell movement is influenced by interactions with neighbouring cells and random motility. Short-range repulsion prevents collisions, while filopodia-mediated long-range adhesion can draw cells closer. We note that filopodia are not drawn to scale as they can reach several cell diameters (Ferrer-Bonet & Ruiz-Trillo, 2017). Based on our experimental observations, the model accounts for increased filopodia-mediated adhesion and elevated random motility with local FBS concentration (see Sections I.1.2-I.1.3 of the Supplementary Material). **b**, A partial differential equation (PDE) is used to describe the dynamics of FBS in the culture medium, accounting for its diffusion and uptake by cells. FBS decay is considered negligible within the timescale of interest. **c,** The full model couples cell dynamics with the continuum description of FBS concentration in the medium.

**Extended Data Figure 2.**
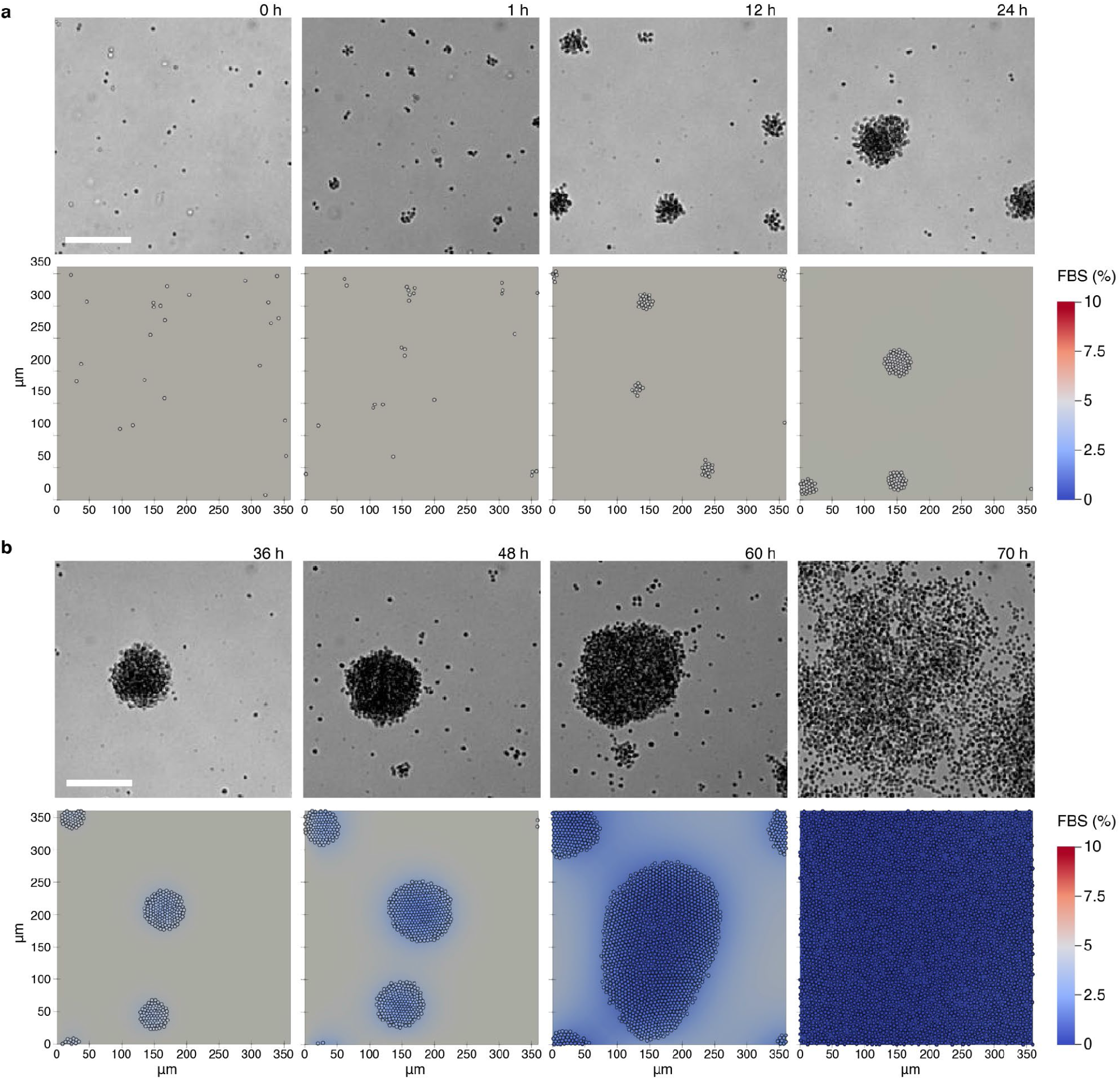
Comparison of *Capsaspora* aggregation between experimental results and model simulations under 5% FBS in a smaller domain. Experimental images (top row) and snapshots of a numerical simulation (bottom row) of chemically induced *Capsaspora* aggregation at **a**, 0-24 and **b**, 36-70 hours after administration of 5% FBS. Both experimental and simulated images cover a 360 x 360 μm domain. Scale bar in experimental images corresponds to 100 μm. In the simulation snapshots, cell colours represent the local FBS concentration, which is also reflected in the background of the medium. The colour bar on the right indicates FBS levels: initial 5% FBS appears grey, while depletion due to metabolic activity of *Capsaspora* shifts colours to blue, with dark blue indicating no FBS. At early time moments of *Capsaspora* aggregation (0-24 h in **a**), no noticeable FBS depletion can be seen. However, lower concentrations of FBS in regions surrounding cell aggregates after 36 h can be observed, which leads to aggregate disassembly at 60 h and complete disaggregation at 70 h. Numerical simulations were performed with sigmoidal Hill response of cell adhesion strength to FBS levels (see Section I of Supplementary Information). All parameter values are as in Extended Data Table 1. For the complete animation of this simulation, see Movie 3.

**Extended Data Figure 3.**
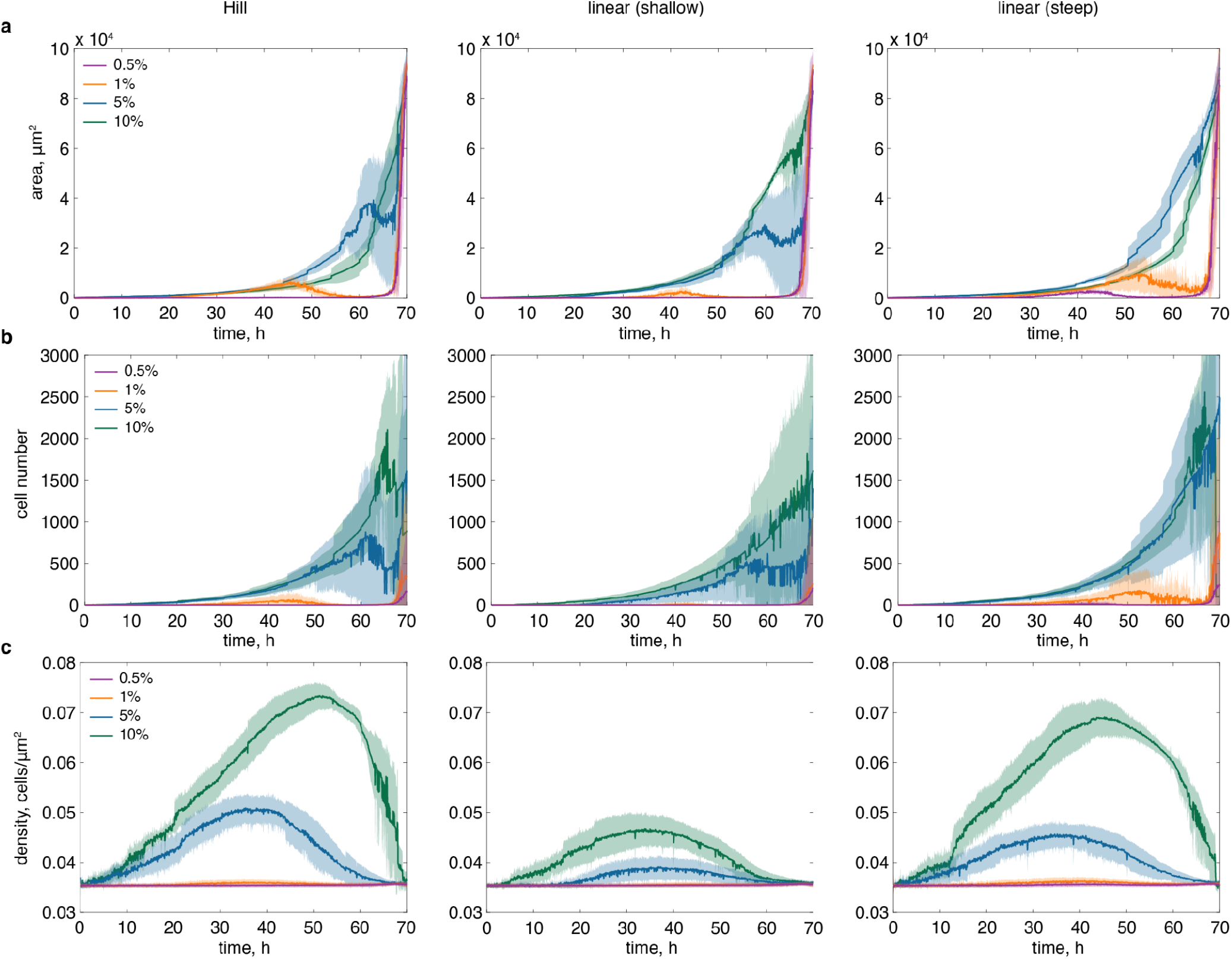
Quantification of the largest multicellular aggregate (by area) in numerical simulations for different initial FBS concentrations. Panels show the temporal evolution of **a**, the mean area of the largest aggregate, **b**, the number of cells within this aggregate, and **c**, the corresponding aggregate density. Solid lines represent the mean values across five numerical simulations for each condition, with shaded regions indicating standard deviation. Colours distinguish simulations with initial FBS levels of 0.5% (purple), 1% (orange), 5% (blue), and 10% (green). The first column corresponds to a threshold-like Hill response of cell adhesion to FBS, the middle column to a moderate linear increase (shallow slope), and the right column to a steep linear response. Animations of individual simulations are available in Movies 6-8 for the Hill, shallow linear, and steep linear responses, respectively. Notably, variance in the metrics increases significantly as aggregate disassembly begins around 55-60 h. Additionally, panel **c** reveals that aggregates are more compact at higher FBS concentrations, explaining why they appear smaller at these levels, as also observed experimentally in (Ros-Rocher et al., 2023). For these simulations, we set the domain size to 360 x 360 μm, and all parameter values are as in Extended Data Table 1.

**Extended Data Figure 4.**
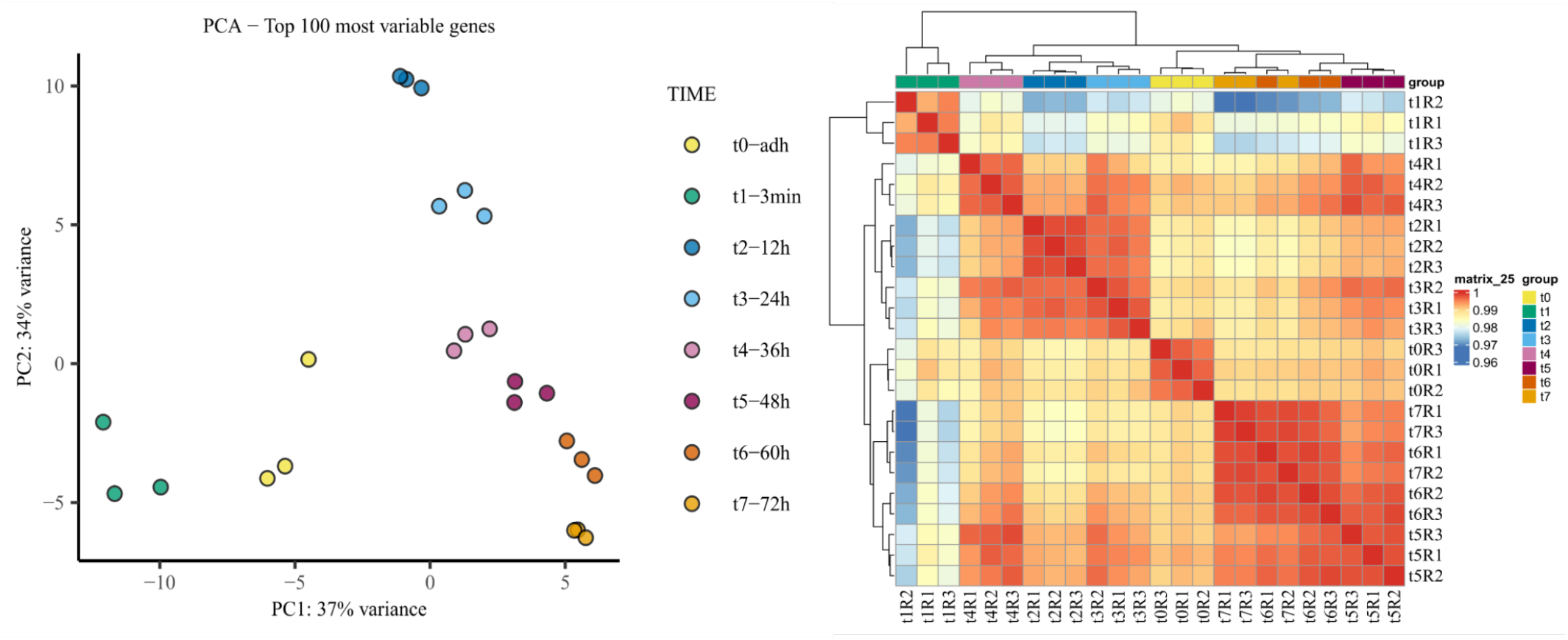
Most variable genes during the aggregation process. **a**, PCA of the top 100 most variable genes at eight different time points of the aggregation process **b**, Heatmap of the top 100 most variable genes forms four clusters based on pairwise correlation representing disaggregation T5, T6 and T7; adherent stage t0; aggregate growth t2, t3, t4; & aggregate induction T1.

**Extended Data Figure 5.**
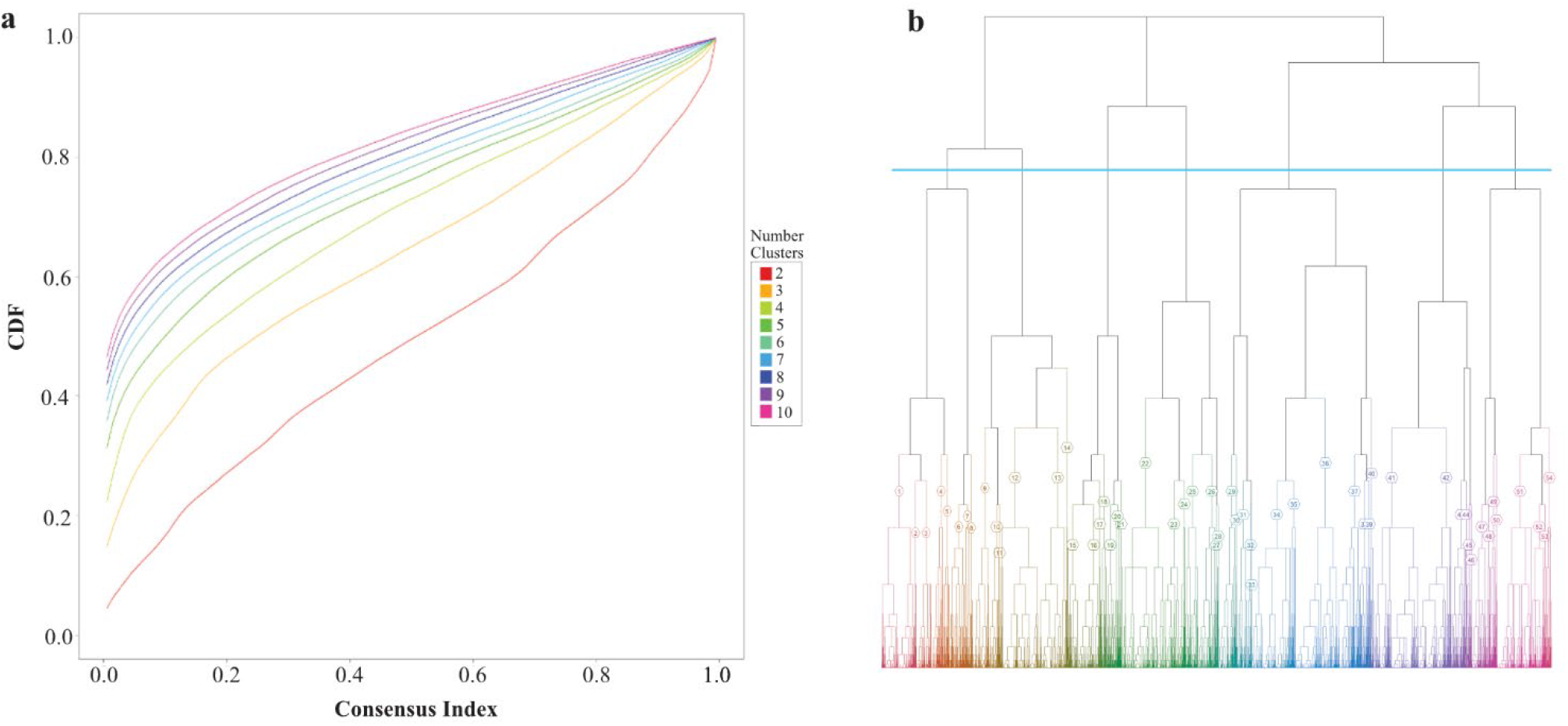
a, Cumulative Distribution Function plot. (**CDF**) calculated for different numbers of gene expression pattern clusters, after 6 clusters (light blue) the curves start to flatten, meaning there is no addition to the total explained variability indicating that 6 clusters are sufficient to capture most of the variability **b**, **DEGpattern dendrogram** of the 54 clusters obtained by hierarchical clustering, the blue line represents the 6 supergroups used for the gene expression pattern analysis (Figure 5)

**Extended Data Figure 6.**
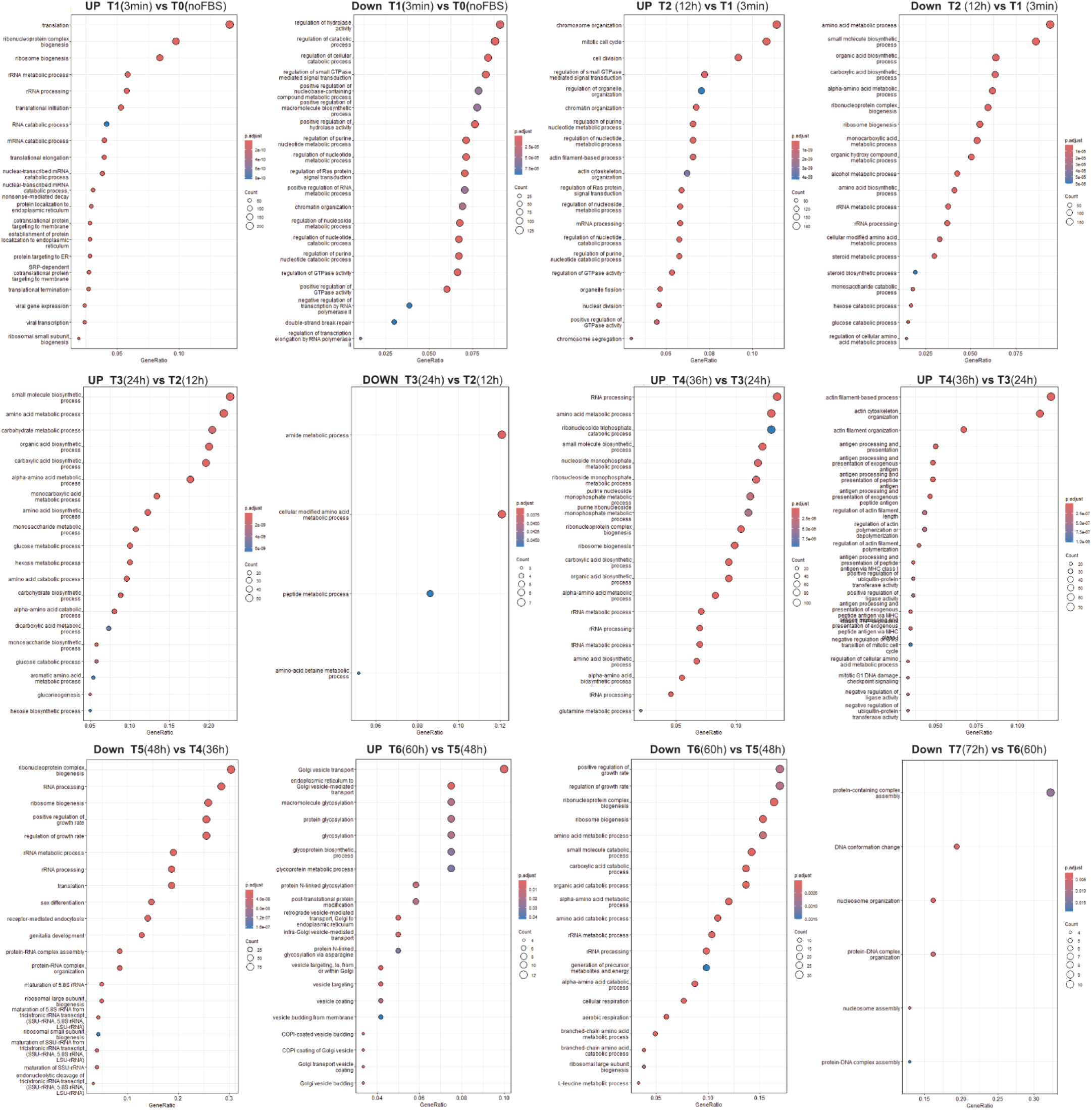
Dotplots of enriched GO terms across the aggregation process. The dotplots represent the top 20 significant GO terms for each of the timepoint pairwise comparisons generated with clusterProfiler, the color of the dots indicates the p-value and the size the number of genes in the GO term, the position indicates the enrichment score, genes with high enrichment score (right position), low adjusted P values (warm colours), and high number of genes associated to the GO (bigger size) are the most significant ones.

**Extended Data Figure 7.**
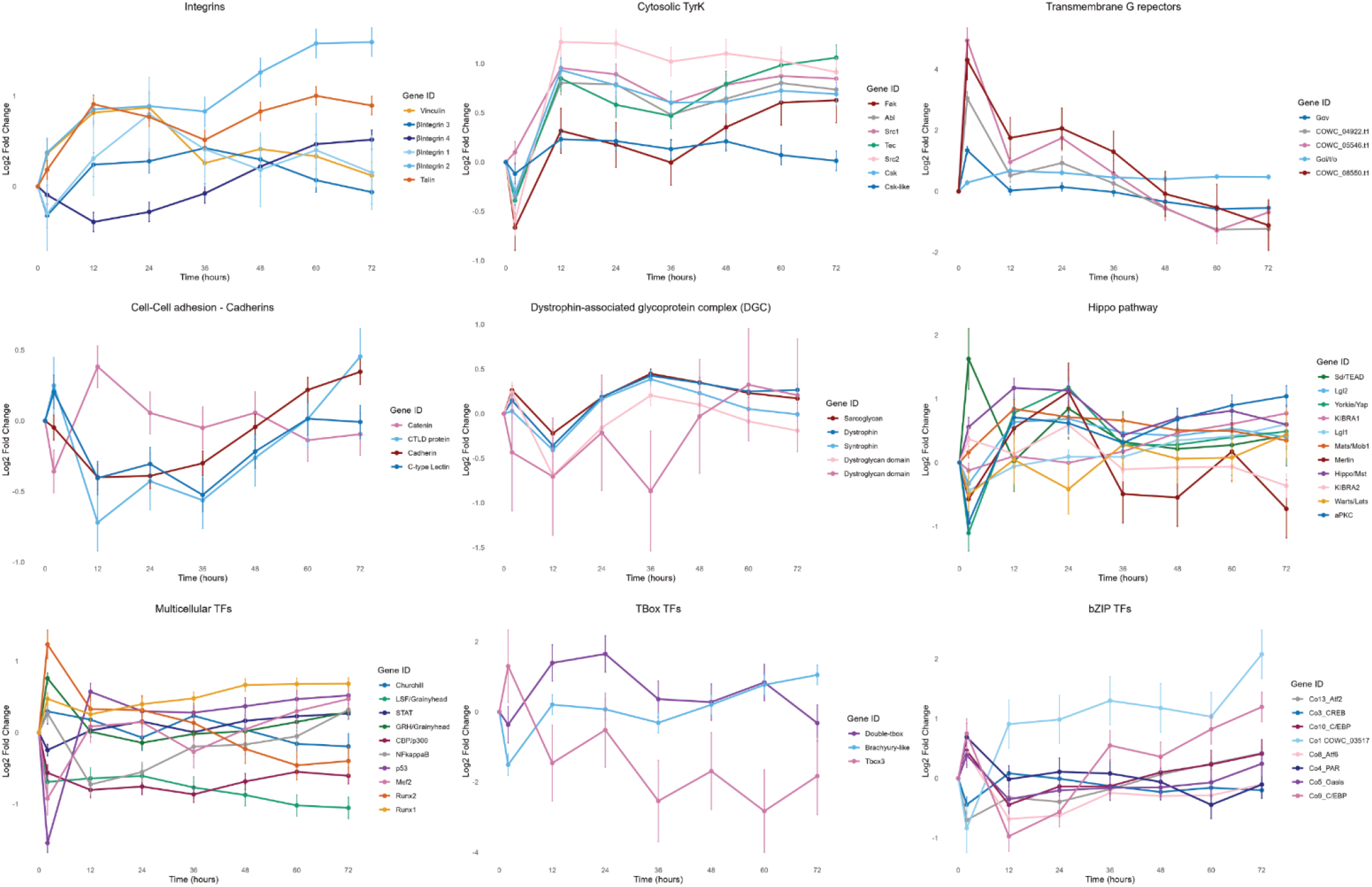
Dynamic expression patterns of genes homologous to metazoan genes related to multicellularity. **a,** Integrins **b,** Cytosolic tyrosine kinases **c,** Transmembrane G receptors **d,** Cadherin **e,** Dystrophin associated glycoprotein complex **f,** Hippo pathway **g,** General transcription factors **h,** TBox transcription factors **i,** bZip transcription factors

**Extended Data Figure 8.**
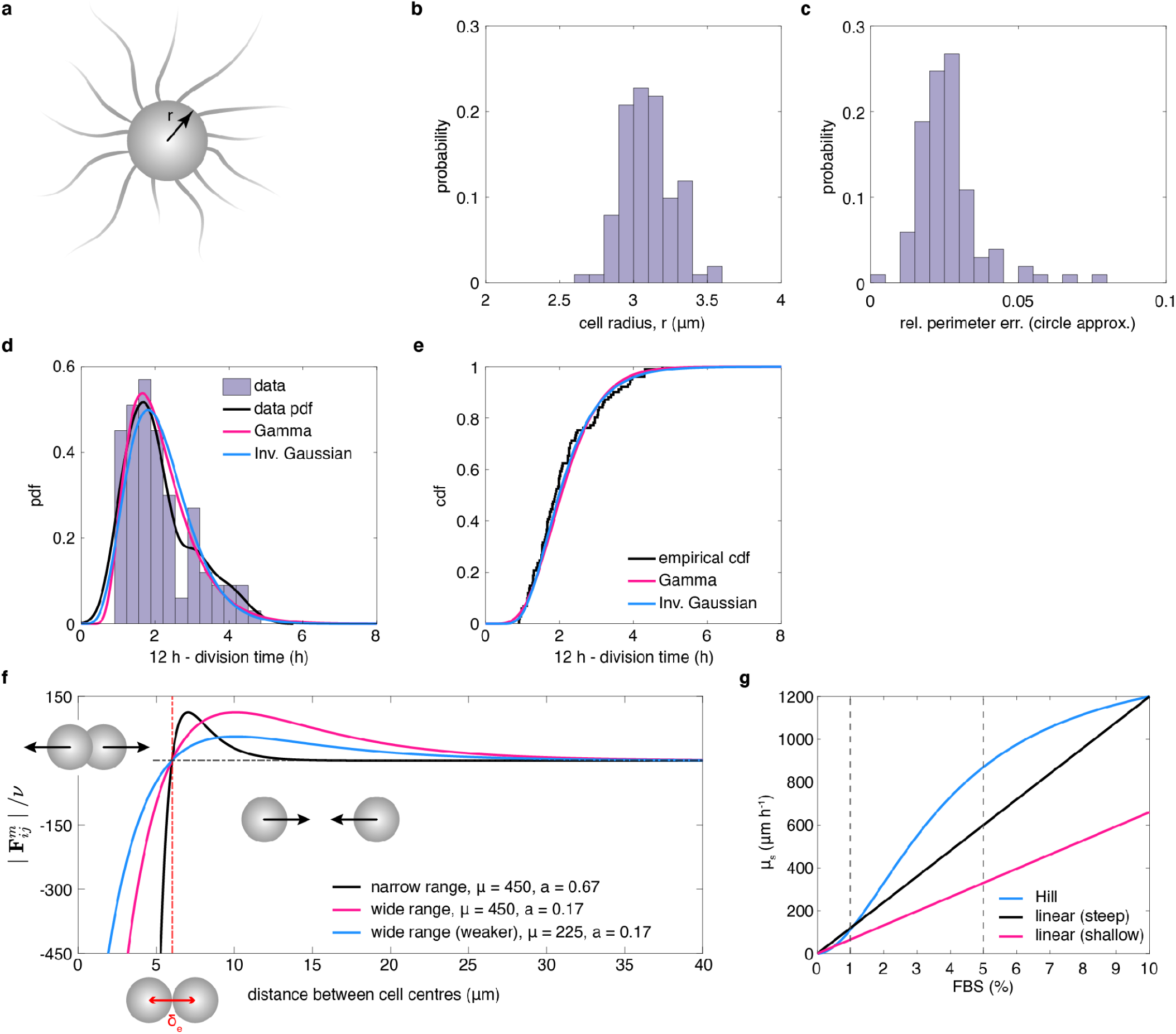
Modelling cell geometry, division dynamics, and adhesion response in *Capsaspora*. **a**, *Capsaspora* cells have spherical bodies with long filopodia (not drawn to scale), which facilitate long-range interactions. In our simulations, we simplify their representation to circular cell bodies. **b**, **c**, Experimental data from (Pérez-Posada et al., 2020) support this circular approximation. **b**, The histogram shows the distribution of cell radii (*r*) inferred from area measurements, with a mean of 3.10 ± 0.17 *μm*. We use *r* = 3 *μm* in simulations. **c**, A comparison of experimental and estimated perimeters (*P*^∗^ = 2*πr*) shows a low mean relative error (< *e_P_* > = 0.0263), validating the circular assumption. **d**, **e**, The cell cycle model was calibrated using *Capsaspora* division time distributions from (Pérez-Posada et al., 2020). **d**, The histogram (purple) shows the distribution of (12 h - division time), fitted to Gamma (magenta) and Inverse Gaussian (blue) distributions. The Gamma shape and scale parameters are *k̃* = 6.30 and *θ̃* = 0.34, while the Inverse Gaussian mean and shape parameters are *μ̃* = 2.15 and *λ̃* = 12.23. **e**, The cumulative density functions (cdfs) confirm the fit of both distributions to experimental data. **f**, The rescaled Morse force magnitude, 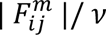, is plotted against cell separation distance *δ_ij_*. The force is repulsive for *δ_ij_* < *δ_e_* (6 *μm*) and attractive for *δi_i_* > *δ_e_*, with zero force at equilibrium. **g**, The FBS-dependent adhesion strength, *μ_s_*(*FBS*), follows three functional forms: a Hill response (blue) and two linear responses with different slopes (magenta and black). Parameter values are listed in Extended Data Table 1.

**Extended Data Figure 9.**
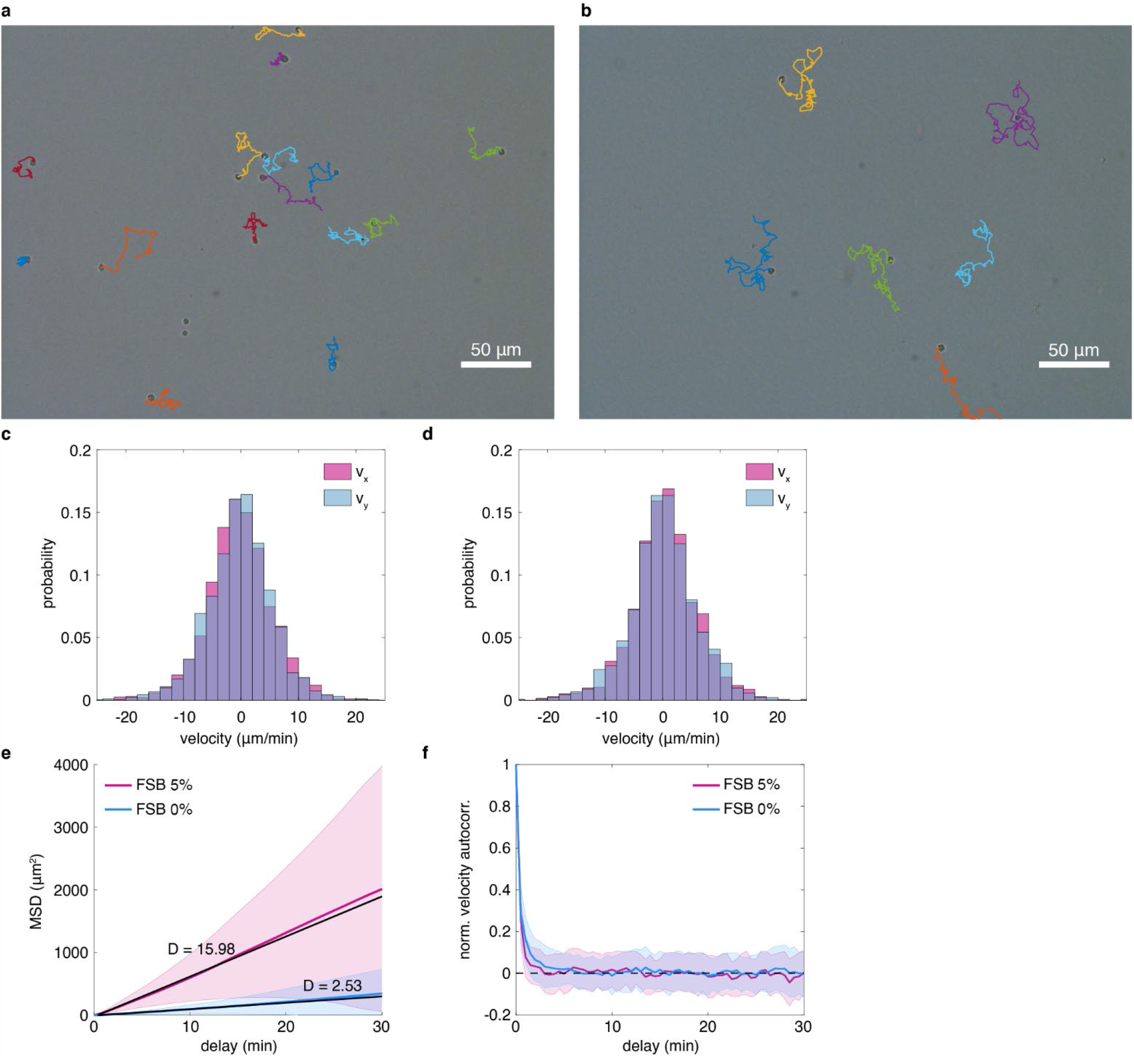
Single-cell motility of *Capsaspora* in different FBS conditions. **a**, **b**, Trajectories of individual cells under **a** 0% FBS and **b** 5% FBS. The images show the initial cell positions, with the full 1-hour trajectories overlaid (colouring is arbitrary). As such, cells are positioned at the start of their respective tracks. Tracks of clustered cells or those present only briefly were excluded from the analysis. **c**, **d**, Velocity distributions of individual cells from **c** *Fpop* and **d** *Cl1* cell lines. The distributions are calculated for 33 tracks in **c** and 14 tracks in **d**. **e**, The ensemble-averaged MSD, extracted from individual cell trajectories, is used to estimate the diffusion coefficient for cells in 0% FBS (blue) and 5% FBS (magenta). Solid lines represent the mean MSD values, *x*^2^(*τ*) > as a function of the delay time *τ* (mins), while the shaded regions indicate the weighted standard deviation. The diffusion coefficient *D* is estimated by fitting a linear function of the form *x*^2^(*τ*) ≥ 2*dDτ*, to the first 20 minutes of the MSD curves. Here, *d* = 2 indicates the spatial dimension. Using this approach, we obtain *D* = 2.53*μm*^2^*min*^−1^for 0% FBS, with a 95%-confidence interval of [2.47,2.59] and a goodness-of-fit, *R*^2^ = 0.995. For 5% FBS, the diffusion coefficient is *D* = 15.98*μm*^2^*min*^−1^, with a 95%-confidence interval of [15.73,16.24] and a goodness-of-fit, *R*^2^ = 0.998. **f**, We also calculated the normalised velocity autocorrelation function, *Z*(*τ*) = *v*(*t* + *τ*), *v*(*t*) >< *v*(*t*), *v*(*t*) >, for individual cell trajectories in 0% FBS (blue) and 5% FBS (magenta). Solid lines show the mean autocorrelation, while the shaded regions represent the weighted standard deviation. The statistics in **e** and **f** are calculated for 78 tracks in 0% FBS and 69 tracks in 5% FBS. See Supplementary Materials, Section I.1.3 for details of parameter estimation.

**Extended Data Table 1.** Descriptions of parameters included in our model and their baseline values used in this work. Details regarding the methods used for parameter estimation are given in the Supplementary Materials, Section I. For simplicity, we omit bars in the notation of the parameters rescaled with cell radius. SI: Supplementary Information.

| Par. | Value | Description |
| --- | --- | --- |
| $r$ | $3\ \mu\text{m}$ | Radius of the *Capsaspora* spherical core. |
| $\delta_e$ | $6\ \mu\text{m}$ | Equilibrium distance for the Morse force for adult *Capsaspora*. |
| $\mu_s$ | $900\ \mu\text{m} \cdot \text{h}^{-1}$ | Rescaled strength of the Morse force ($\mu_s = \hat{\mu}_s/\nu$, where $\nu$ denotes the drag coefficient) from Eq (7) in SI. |
| $a_s$ | $0.67\ \mu\text{m}^{-1}$ | Parameter controlling the range of interaction for short -range repulsion in Eq (7) in SI. |
| $a_l$ | $0.17\ \mu\text{m}^{-1}$ | Parameter controlling the range of interaction for long -range adhesion mediated by filopodia in Eq (7) in SI. |
| $\mu_0$ | $1412.1\ \mu\text{m} \cdot \text{h}^{-1}$ | Maximum value of the rescaled adhesion strength of the Morse force for the Hill response to FBS (see blue curve in Extended Data Figure 8g). |
| $f_0$ | 3.8494 % | FBS concentration producing half of the maximum adhesion strength in the Hill response to FBS (see blue curve in Extended Data Figure 8g). |
| $n_0$ | 1.8153 | Hill coefficient in the Hill response to FBS of *Capsaspora* adhesion strength (see blue curve in Extended Data Figure 8g). |
| $k_\mu^1$ | $66\ \mu\text{m} \cdot \text{h}^{-1}$ | Slope parameter in the linear response (with shallow slope) to FBS of *Capsaspora* adhesion strength (see magenta curve in Extended Data Figure 8g). |
| $k_\mu^2$ | $120\ \mu\text{m} \cdot \text{h}^{-1}$ | Slope parameter in the linear response (with steep slope) to FBS of *Capsaspora* adhesion strength (see black curve in Extended Data Figure 8g). |
| $k$ | 2.6906 | Slope parameter in the linear dependency of the *Capsaspora* diffusion coefficient on FBS concentration (Eq (8) in SI). |
| $b$ | 2.53 | Intercept parameter in the linear dependency of the *Capsaspora* diffusion coefficient on FBS concentration (Eq (8) in SI). |
| $D_{fbs}$ | $675\ \mu\text{m}^2 \cdot \text{h}^{-1}$ | FBS diffusion coefficient. |
| $k_u$ | $20.7\ \mu\text{m}^2 \cdot \text{cell}^{-1} \text{h}^{-1}$ | Rate of FBS consumption per cell per area of the finite element model (FEM) element (in our simulations, a right triangle with sides equal to $14.4\ \mu\text{m}$). |
| $L$ | 360 or 1080 $\mu\text{m}$ | Size of the square domain used in the model simulations. |
| $N_0$ | 24 or 216 cells | Initial cell number scattered across the simulation domain at $t = 0$ for domain with side of $360\ \mu\text{m}$ and $1080\ \mu\text{m}$, respectively. |
| $\tilde{\mu}$ | 2.1530 | The mean parameter of the Inverse Gaussian distribution described by the probability density function from Eq (3) in SI which provided the best fit for the experimental data of the quantity [12 h - division time (h)]. |
| $\tilde{\lambda}$ | 12.2344 | The shape parameter of the Inverse Gaussian distribution described by the probability density function from Eq (3) in SI which provided the best fit for the experimental data of the quantity [12 h - division time (h)]. |
| Parameter values after rescaling the space with the cell radius. |  |  |
| $r$ | 1 | Radius of the *Capsaspora* spherical core. |
| $\mu_s$ | $300\ \text{h}^{-1}$ | Rescaled strength of the Morse force. |
| $a_s$ | 2.0 | Parameter controlling the range of interaction for short -range repulsion. |
| $a_l$ | 0.5 | Parameter controlling the range of interaction for long -range adhesion via filopodia. |
| $\mu_0$ | $470.7\ \text{h}^{-1}$ | Maximum value of the rescaled adhesion strength of the Morse force for the Hill response to FBS (see blue curve in Extended Data Figure 8g). |
| $k_\mu^1$ | $22\ \text{h}^{-1}$ | Slope parameter in the linear response (with shallow slope) to FBS of *Capsaspora* adhesion strength (see magenta curve in Extended Data Figure 8g). |
| $k_\mu^2$ | $40\ \text{h}^{-1}$ | Slope parameter in the linear response (with steep slope) to FBS of *Capsaspora* adhesion strength (see black curve in Extended Data Figure 8g). |
| $k$ | 0.2990 | Slope parameter in the linear dependency of the *Capsaspora* diffusion coefficient on FBS concentration. |
| $b$ | 0.2811 | Intercept parameter in the linear dependency of the *Capsaspora* diffusion coefficient on FBS concentration. |
| $D_{fbs}$ | $75\ \text{h}^{-1}$ | FBS diffusion coefficient. |
| $k_u$ | $2.3\ \text{cell}^{-1} \text{h}^{-1}$ | Rate of FBS consumption per cell per area of the FEM element (in our simulations, a right triangle with sides equal to 4.8, dimensionless). |
| $L$ | 120 or 360 | Size of the square domain used in the model simulations. |

