## Supplementary information for "An experimentally tractable framework for quantitative and integrative analysis of aggregation in the origin of animals"

[*] Corresponding authors:

### I Detailed model description

We present a two-dimensional (2D) model of *Capsaspora* aggregation, regulated by externally supplied FBS (Fetal Bovine Serum). The framework comprises two key components: an agent-based model for *Capsaspora* cells and a continuum description of the FBS dynamics. These two submodels are coupled by assuming that cell behaviour is influenced by the local FBS concentration while FBS is consumed by the cells. The model is implemented using *Chaste*, an open-source library for simulating biological systems (Mirams et al., 2013). We also emphasise that, wherever possible, model parameters were derived from available experimental data to ensure our model accurately captures the observed dynamics of *Capsaspora* aggregation. Below, we provide a detailed description of each modelling component.

#### I.1 Cell model

To model the aggregation of *Capsaspora*, we develop a cell-centred agent-based model that simulates FBS-dependent cell motility and mechanical cell-cell interactions, and cell proliferation. In this framework (see, for example, (Bull et al., 2020)), cells are represented as spheres (or circles in 2D) with radius $r$ (see Extended Data Figure 8a-c) and move in an off-lattice manner, influenced by forces acting upon them.

The assumption of circular cell geometry is supported by experimental measurements of cell morphology at division reported in (Pérez-Posada et al., 2020) (see Extended Data Figure 8b-c). While *Capsaspora* cells possess long, thin filopodia that can extend up to $2-4$ cell diameters in length (Ferrer-Bonet & Ruiz-Trillo, 2017), we simplify our model by representing only the circular cell body. While we assume that filopodia play a role in facilitating long-range cell-cell interactions (see also Section I.1.2), for simplicity, we will represent only the circular cell body in our simulations.

The forces acting on cells result from a combination of mechanical interactions with neighbouring cells and random motility. Accordingly, the position of the $i$th cell evolves over time following the equation:

$$\frac{\mathrm{d}\boldsymbol{x}_i}{\mathrm{d}t} = \frac{1}{\nu}\boldsymbol{F}_i^m + \boldsymbol{F}_i^d. \tag{1}$$

Here, $\boldsymbol{x}_i$ denotes the position of the centre of cell $i$, $\boldsymbol{F}_i^m$ represents mechanical forces from cell-cell interactions and $\boldsymbol{F}_i^d$ corresponds to random cell motility. The parameter $\nu$ is the drag coefficient. Since cell size is significantly larger than the molecules in the surrounding liquid medium, viscous effects are predominant, and inertial forces can be assumed negligible. To complete Eq (1), the initial position of each cell must be specified as $\boldsymbol{x}_i(t=0) = \boldsymbol{x}_i^0$. We typically use random initial conditions representing initial cell scattering across a two-dimensional square domain (see also Section I.1.4). For simplicity, we also assume that the cell radius, $r$, is uniform across the *Capsaspora* population.

We provide a detailed description of the implementation of mechanical cell-interaction forces in Section I.1.2. Random cell motility is modelled as Brownian motion, as follows

$$\boldsymbol{F}_i^d = \boldsymbol{\xi}_i(t), \tag{2}$$

where $\boldsymbol{\xi}_i(t)$ is a Gaussian process with zero mean, $\langle \boldsymbol{\xi}_i(t) \rangle = 0$ for any $t$, and the covariance given by $\langle \boldsymbol{\xi}_i(t), \boldsymbol{\xi}_i(t') \rangle = 2D_i\delta(t-t')$ for any $t$, $t'$. Here, $D_i$ is the diffusion coefficient that characterises the motility of cell $i$, and $\delta(t)$ is the Dirac delta function. *Capsaspora* motility is upregulated in response to externally supplied FBS. Thus, the value of $D_i$ depends on the FBS concentration perceived by cells in their microenvironment. In Section I.1.3, we estimate the FBS-dependent diffusion coefficient $D_i$ from single-cell tracking data and justify our assumption that *Capsaspora* random movement can be modelled as Brownian motion.

Beyond migration, our model also incorporates *Capsaspora* proliferation. A detailed explanation of the cell cycle model is provided in Section I.1.1.

##### I.1.1 Cell cycle

We use experimental data from (Pérez-Posada et al., 2020) (see Fig 2C therein) to characterise the distribution of *Capsaspora* division times. This dataset, which includes $101$ division events, shows that cell proliferation occurred between $7$ and $11$ hours, with a mean division time of $9.87$ hours and a standard deviation of $0.91$ hours. To model the distribution of *Capsaspora* cell cycle lengths, we fitted a probability distribution to the observed division times. To simplify the fitting process, we transformed the division time data to $[12h -$ division time (h)$]$. Extended Data Figure 8d illustrates this transformation and the best fits for the Gamma and Inverse Gaussian distributions. These two distributions were chosen for their ability to capture the heavy tail observed in the experimental data (see Extended Data Figure 8d). In Extended Data Figure 8e, we present the cumulative distribution functions for both the experimental data and the fitted distributions. Both the Gamma and Inverse Gaussian distributions provide a reasonable fit to the data. However, a goodness-of-fit test indicates that the Inverse Gaussian distribution offers a slightly better fit, with a $p$-value of $p = 0.184$ compared to $p = 0.1059$ for the Gamma distribution. Therefore, we use the Inverse Gaussian distribution in our simulations. Its probability density function is given by the following expression:

$$f_{IG}(x; \mu, \lambda) = \sqrt{\frac{\lambda}{2\pi x^3}} \exp\left[-\frac{\lambda (x-\mu)^2}{2\mu^2 x}\right], \tag{3}$$

where $\mu$ is the mean and $\lambda$ is the shape parameter.

Pérez-Posada et al. (2020) also confirmed that *Capsaspora* division is symmetric (see Fig 2D therein), with each daughter cell getting approximately equal volume of the parent cell. Based on this, we implement our cell cycle model as follows. Each cell $i$ is characterised by its birthtime, $t_i^b$, and the time of its next division, $t_i^d$. During our numerical simulations, we perform the following steps.

- $t = 0$ At the start of each simulation (simulation time $t = 0$), each cell agent is assigned a birth time in the past as $t_i^b = -U_{[0,1]} \cdot 9.87$ h, where $9.87$ is the mean division time from (Pérez-Posada et al., 2020), and $U_{[0,1]}$ is a uniform distribution on the interval $[0, 1]$. The next division time is set to $t_i^d = 12 - \zeta$, where $\zeta$ is drawn from the Inverse Gaussian distribution $\mathrm{IG}(\tilde{\mu}, \tilde{\lambda})$ (see Extended Data Figure 8d-e).
- $t = t_i^d$ As the simulation progresses and reaches the division time for the $i$-th cell, the cell divides into two daughter cells with indices $\bar{i}_1$ and $\bar{i}_2$, each with a radius of $r/2$. The centres of the daughter cells are positioned $r/2$ apart from the parent cell centre, $\boldsymbol{x}_i$, along a randomised direction. The birth times are set to the current simulation time as $t_{\bar{i}1}^b = t$ and $t_{\bar{i}2}^b = t$. The next division times are set to $t_{\bar{i}1}^d = t + (12 - \zeta_1)$ and $t_{\bar{i}_2}^d = t + (12 - \zeta_2)$, where $\zeta_1$ and $\zeta_2$ are drawn from $\mathrm{IG}(\tilde{\mu}, \tilde{\lambda})$.
- $t = t_i^d + 2$ Over the following $2$ hours, the newly formed daughter cells grow linearly until their radii reach $r$. This allows for a more realistic implementation of cell division and also ensures no numerical instabilities occur during simulations.
- $t = t_{final}$ The steps above are repeated for all cells throughout the simulation until the final simulation time is reached.

##### I.1.2 Cell interaction forces

As discussed earlier, we assume that cell position is influenced by random motility and cell interaction forces, represented by the terms $\boldsymbol{F}_i^d$ and $\boldsymbol{F}_i^m$ in Eq (1), respectively. Regarding the cell-cell interaction force, $\boldsymbol{F}_i^m$, we assume that *Capsaspora* cells exhibit repulsive interactions at short distances (to prevent collisions) and adhesive interactions at longer distances, where filopodia can become entangled and pull cells together into clusters. The interaction force is also assumed to weaken with increasing separation distance and become negligible beyond the typical filopodium length. In this study, we model this behaviour using the Morse potential. The Morse potential, $V(\delta_{ij})$, describes the interaction energy between two cells separated by a distance $\delta_{ij}$, and is given by the following expression:

$$V(\delta_{ij}) = D_e(1 - \exp(-a(\delta_{ij} - \delta_e)), \tag{4}$$

where $D_e$ determines the depth of the potential well, controlling the strength of the interaction, and $a$ regulates the width of the well. The term $\delta_{ij} = \|\boldsymbol{x}_i - \boldsymbol{x}_j\|$ represents the distance between cells $i$ and $j$, and $\delta_e$ is the equilibrium separation distance.

In the case of our model with spherical cells, the equilibrium distance between two interacting cells is equal to the sum of their radii, i.e. $\delta_e = r_i + r_j$. For fully grown (adult) cells, this quantity is equal to $\delta_e = 2r$. Shorter distances between cells $i$ and $j$ ($\delta_{ij} < \delta_e$) will result in repulsion. Conversely, for separation distances larger than the equilibrium distance ($\delta_{ij} > \delta_e$), Eq (4) will give rise to attracting adhesion forces between cells. Increasing $D_e$ results in stronger interaction forces between cells, while decreasing $a$ extends the range of their interaction. The interaction force between cells $i$ and $j$ is then given by $\boldsymbol{F}_{ij}^m = (-\nabla_x V(\delta_{ij}), -\nabla_y V(\delta_{ij}))^T$, where the gradients of the Morse potential determine the force components in the $x$ and $y$ directions:

$$\boldsymbol{F}_{ij}^m = \widehat{\mu} \left[\exp(-a(\delta_{ij} - \delta_e)) - \exp(-2a(\delta_{ij} - \delta_e))\right] \frac{1}{\delta_{ij}} (\boldsymbol{x}_i - \boldsymbol{x}_j), \tag{5}$$

where we defined $\widehat{\mu} = 2aD_e$. Extended Data Figure 8f shows how the shape of the Morse force varies for three different combinations of the parameters $\widehat{\mu}$ and $a$, which control the strength and range of the interaction, respectively. Finally, the total force experienced by cell $i$ is obtained by summing $\boldsymbol{F}_{ij}^m$ over all interacting cells in the system:

$$\boldsymbol{F}_i^m = \sum_j \boldsymbol{F}_{ij}^m. \tag{6}$$

The numerical implementation of Eq (6) can be simplified by summing only over cells within the maximum interaction range, $\delta_{max}$, of cell $i$.

Although the interaction force described by Eqs (5)-(6) accounts for both short-range repulsion and long-range adhesion, we will discuss these two processes separately, as they are driven by distinct biological mechanisms.

**Short-range repulsion** Short-range repulsion arises from the viscoelastic properties of cells when they are in close proximity ($\delta_{ij} < \delta_e$). These repulsive forces prevent cells from overlapping or invading each other's space, as two spherical cells cannot occupy the same volume. Such interactions are purely mechanical, and we assume them to be independent of external factors like local FBS concentration. We denote by $\widehat{\mu}_s$ and $a_s$ the parameters of the Morse force $\boldsymbol{F}_i^m$ for $\delta_{ij} \leq \delta_e$, which are considered to be fixed during numerical simulations.

**Long-range adhesion** We assume that *Capsaspora* utilise their long filopodia to aggregate and can adhere to each other across longer distances by using their filopodia (Ros-Rocher et al., 2023). Given that *Capsaspora* do not form aggregates in FBS-free media but begin aggregating when FBS is supplied (Ros-Rocher et al., 2023), we hypothesise that the magnitude of adhesion forces depends on the local FBS concentration sensed by the cells. Thus, we assume that the parameter regulating the strength of adhesive interactions is an increasing function of the FBS concentration, $\widehat{\mu}_l = \widehat{\mu}_l(\text{FBS})$. Specific functional forms of $\widehat{\mu}_l(\text{FBS})$ are discussed below. Conversely, the parameter $a_l$, which controls the range of interaction between cells, is assumed constant. Its value was estimated in our simulations to ensure non-zero adhesion forces between cells separated by distances typical for their filopodium length, which can span $3-4$ cell diameters, approximately corresponding to $20-25\ \mu m$.

Combining short-range repulsion and longer-range adhesion, we obtain the following functional form for the cell interaction force that we employ during our numerical simulations:

$$\frac{1}{\nu}\boldsymbol{F}_{ij}^{m} = \begin{cases} \mu_s \left[\exp\left(-a_s(\delta_{ij}-\delta_e)\right) - \exp\left(-2a_s(\delta_{ij}-\delta_e)\right)\right] \frac{1}{\delta_{ij}} \left(\boldsymbol{x}_i - \boldsymbol{x}_j\right), & \text{if } \delta_{ij} \leq \delta_e, \\ \mu_l(\mathrm{FBS}_{ij}) \left[\exp\left(-a_l(\delta_{ij}-\delta_e)\right) - \exp\left(-2a_l(\delta_{ij}-\delta_e)\right)\right] \frac{1}{\delta_{ij}} \left(\boldsymbol{x}_i - \boldsymbol{x}_j\right), & \text{otherwise.} \end{cases} \tag{7}$$

Here, we directly rescaled the interaction force with the drag coefficient so that $\mu_s = \widehat{\mu}_s/\nu$ and $\mu_l = \widehat{\mu}_l/\nu$. The equilibrium distance between cells, $\delta_e$, is calculated as a sum of radii for the interacting cells $i$ and $j$ (which is equal to $\delta_e = 2r$ for fully grown cells; see also Section I.1.1). The local FBS concentration sensed by cells $i$ and $j$ is calculated as $\mathrm{FBS}_{ij} = 0.5\,(\mathrm{FBS}_i + \mathrm{FBS}_j)$, where $\mathrm{FBS}_i$ and $\mathrm{FBS}_j$ are local FBS concentrations sensed by cells $i$ and $j$, respectively. The total force experienced by cell $i$ can then be obtained by summing over all neighbouring cells $j$ as shown in Eq (6). We report the baseline parameter values in Extended Data Table 1 (upper part).

In this work, we explore three alternative functional forms for the FBS-upregulated adhesion strength, $\mu_l(\mathrm{FBS})$. As demonstrated by (Ros-Rocher et al., 2023), *Capsaspora* metabolise FBS components, and the resulting metabolic products localise along their filopodia, enhancing cell-cell connections. This supports our assumption that the strength of adhesive forces increases with FBS concentration, which can be interpreted as an increase in the 'stickiness' of *Capsaspora* filopodia at higher FBS levels. Moreover, the results from (Ros-Rocher et al., 2023) on mean aggregate size in response to increasing FBS concentrations suggest a highly non-linear reaction to this stimulus (see Fig. 2D therein). Specifically, their experiments showed that cells did not form aggregates at FBS levels below $1\%$, formed small, low-density aggregates at FBS concentrations at $1.0-1.5\%$, and created larger, more defined aggregates at higher concentrations. This motivated us to model $\mu_l(\mathrm{FBS})$ using a non-linear, sigmoidal response captured by a Hill function, i.e. $\mu_l(\mathrm{FBS}) = H(\mathrm{FBS}; \mu_0, f_0, n_0) = \mu_0/\left[1 + (f_0/\mathrm{FBS})^{n_0}\right]$, with $\mu_0$, $f_0$ and $n_0$ being the parameters defining the shape of the Hill function. Such a response function enables us to capture better the initial sharp increase in adhesion strength around $1\%$ FBS, followed by a slower saturation at higher concentrations (blue curve in Extended Data Figure 8g). As an alternative, we also considered a simple linear response, $\mu_l(\mathrm{FBS}) = k_\mu \mathrm{FBS}$, characterised by two distinct slopes: $k_\mu^1$ for a shallower response and $k_\mu^2$ for a steeper response (depicted in magenta and black in Extended Data Figure 8g, respectively). The slope of the shallower linear response was chosen to align with the results observed for the Hill function at lower FBS concentrations while diverging at higher levels. Conversely, the steeper linear response was designed to produce distinct results at low FBS concentrations compared to the sigmoidal response while converging to similar responses at higher FBS levels.

##### I.1.3 Estimation of cell motility from single cell tracking data

In order to estimate individual cell motility, we conducted experiments using *Capsaspora* cells at low cell density on regular surface plates. In all experimental conditions, cells were counted and seeded at the specified densities, incubated at $23°\mathrm{C}$ for 16–20 hours, and then observed under a microscope at 20x magnification. Images were captured at 30-second intervals over a 60-minute period.

To analyse the cell trajectories, we implemented a tracking pipeline using the TrackMate plug-in in ImageJ. We manually reviewed the resulting tracks, excluding any cells that interacted with others or formed clusters. For cells that began moving individually but later formed clusters, only the portion of the trajectory where the cell moved independently was analysed. Extended Data Figures 9a and b illustrate typical cell trajectories over a 1-hour period for 0% and 5% FBS, respectively. An initial visual inspection of these tracks suggests that cells stimulated with FBS exhibit increased motility compared to those grown under no-FBS conditions.

We worked with two *Capsaspora* cell lines: the original population, referred to as *Fpop*, and a clonal population, *Cl1*. First, we assessed whether the two cell lines exhibited similar motility. Given that cell motility increases with FBS stimulation, we conducted this analysis under 5% FBS. We tracked 33 individual *Fpop* cells and 14 *Cl1* cells. The velocity distributions for each cell line are shown in Extended Data Figures 9c and d, for *Fpop* and *Cl1*, respectively. Our analysis shows that under 5% FBS stimulation, the velocity for both cell lines typically does not exceed $25\ \mu m/min$. Additionally, consistent with our hypothesis that *Capsaspora* movement can be described as unbiased Brownian motion, the velocity distributions for both *Fpop* and *Cl1* cell lines have the shape of a symmetric Gaussian centred at $0$. Unfortunately, the number of tracks was insufficient for a reliable statistical comparison between the velocity distributions of the two cell lines. However, since the cells do not show any drastically different behaviour (see Extended Data Figure 9), we assume the motility of *Fpop* and *Cl1* to be similar. Therefore, we combined the tracks from both cell lines to estimate the diffusion coefficient.

We obtained 78 and 69 cell tracks for the 0% and 5% FBS conditions, respectively. To estimate the diffusion coefficient $D$ from Eq (2), we first calculated the mean square displacement (MSD), $\langle \boldsymbol{x}^2(\tau)\rangle = \langle \boldsymbol{x}(t+\tau)-\boldsymbol{x}(t)\rangle^2$, which represents the average squared distance a cell travels over a given delay time, $\tau$. The diffusion coefficient is then estimated by fitting a linear function to the ensemble-averaged MSD across all delay times and tracks. For Brownian motion, the MSD follows the relationship $\langle \boldsymbol{x}^2(\tau)\rangle = 2dD\tau$, where $d$ is the spatial dimension ($d = 2$ for two-dimensional cell motion). Extended Data Figure 9e shows the ensemble MSD for cell tracks in both 0% and 5% FBS conditions. It is clear that cell motility is significantly higher in 5% FBS compared to the no-FBS condition.

To estimate $D$, we fitted a linear function to the ensemble MSD data for the first 20 minutes, as statistical noise increases for larger delay times (see Extended Data Figure 9e). The resulting diffusion coefficients are $D = 2.53\ \mu m^2/min$ for 0% FBS and $D = 15.98\ \mu m^2/min$ for 5% FBS. These results indicate that *Capsaspora* motility in 5% FBS is more than six times greater than under no-FBS conditions. Based on these results, in our simulations, we employ a linear function to describe the dependence of $D$ on the concentration of FBS for concentrations up to $5\%$. For higher concentrations, we assume that metabolic activation remains similar to that at $5\%$, and thus, we employ a constant diffusion coefficient of $D = 15.98\ \mu m^2/min$ for FBS concentrations above $5\%$. This is implemented as follows

$$D_i = D(\mathrm{FBS}_i) = \begin{cases} k \cdot \mathrm{FBS}_i + b, & \text{if } \mathrm{FBS}_i \leq 5, \\ 15.98, & \text{otherwise}, \end{cases} \tag{8}$$

with $k = 2.6906$, $b = 2.53$ and $\mathrm{FBS}_i$ is the FBS concentration sensed by cell $i$. It is straightforward to check that $D(0) = 2.53\ \mu m^2/min$ and $D(5) = 15.98\ \mu m^2/min$ as estimated from experimental data.

By examining Extended Data Figure 9e, it can be argued that anomalous diffusion could better describe *Capsaspora* motility. In the case of anomalous diffusion, the ensemble-averaged MSD scales as $\langle \boldsymbol{x}^2(\tau)\rangle \propto \tau^\alpha$, where $\alpha \neq 1$. Specifically, the data in Extended Data Figure 9e suggests $\alpha > 1$, indicating that *Capsaspora* cells exhibit short-term persistence in their movement direction. To explore this further, we calculated the normalised velocity autocorrelation, shown in Extended Data Figure 9f. This metric assesses whether cells 'remember' their previous direction of movement. For pure Brownian motion, the normalised velocity autocorrelation is 1 at zero delay ($\tau = 0$) and zero for all other delay times. As seen in Extended Data Figure 9f, velocity autocorrelation remains positive for delays $\tau < 3$ minutes and fluctuates around zero for longer times. This suggests that *Capsaspora* 'memory' effects persist only on the timescale of a few minutes, which can be attributed to their mode of migration by pulling on their filopodia (which can occur on the scale of a few minutes). Since we are interested in modelling *Capsaspora* aggregation over a much longer timescale (60-70 hours in our experiments), we consider Brownian motion implemented in our model to be a reasonable approximation of *Capsaspora* movement.

##### I.1.4 Domain size and initial cell distribution

Cell migration is modelled on a 2D domain of width $L$, $\Omega = [0, L]^2$, which provides a reasonable approximation of *Capsaspora* aggregation. In experiments, most cells tend to sediment to the bottom of the seeding plates and form aggregates at a similar height within the culture medium. We impose periodic boundary conditions in both the $x$ and $y$ directions (representing a torus), meaning that if a cell exits the domain through one boundary, it re-enters from the opposite side. We choose this boundary condition to eliminate potential boundary effects.

At the initial time moment, we assume that $N_0$ cells are randomly distributed across the simulation domain. The values of $N_0$ and $L$ are listed in Extended Data Table 1.

#### I.2 FBS dynamics

We assume that *Capsaspora* behaviour, such as motility and adhesion strength (i.e. 'stickiness' of their filopodia), depends on the local FBS concentration. Therefore, to properly capture cell behaviour, we incorporate in our model the FBS dynamics in the culture medium. We use a partial differential equation (PDE) to describe FBS diffusion and uptake by *Capsaspora* metabolic activity:

$$\frac{\partial \mathrm{FBS}}{\partial t} = D_{fbs} \nabla^2 \mathrm{FBS} - k_u \rho(\boldsymbol{x}) \mathrm{FBS}. \tag{9}$$

Here, $D_{fbs}$ is the diffusion coefficient of FBS, $k_u$ is the FBS uptake rate by cells and $\rho(\boldsymbol{x})$ is the local cell density. We use *Chaste* implementation of the finite elements method to solve parabolic PDEs such as Eq (9). This method assumes the discretisation of the domain on smaller subdomains called finite elements. Thus, we calculate the local cell density, $\rho(\boldsymbol{x})$, as the number of cell centres falling within each element divided by the element area. This is implemented within standard *Chaste* routines for PDEs. We assume that FBS is administered at the initial time moment at a uniform concentration, i.e. $\mathrm{FBS}(t = 0, \boldsymbol{x}) = \mathrm{FBS}_0$ for all $\boldsymbol{x} \in \Omega$. The initial FBS concentration is set up to match the experimental conditions. In agreement with our cell model, we use periodic boundary conditions for our PDE description of FBS dynamics ($\mathrm{FBS}(t, x = 0) = \mathrm{FBS}(t, x = L)$ and $\mathrm{FBS}(t, y = 0) = \mathrm{FBS}(t, y = L)$).

We assume slower effective diffusion of FBS compared to its individual components to capture the crowding effects of diffusion in a medium with multicellular clusters ($D_{fbs}$). The FBS uptake parameter, $k_u$, was fit to match the disaggregation time in experimental videos when *Capsaspora* cells consume all available FBS and scatter as they cannot form clusters without additionally supplied FBS.

#### I.3 Rescaling space with the cell radius

For simplicity of numerical implementation of our model in *Chaste*, we rescaled the space variable with *Capsaspora* cell radius as $\overline{\boldsymbol{x}} = \boldsymbol{x}/r$. Thus, we reformulated our model equations in terms of the dimensionless space as follows:

$$\frac{\mathrm{d}\overline{\boldsymbol{x}}_i}{\mathrm{d}t} = \overline{\boldsymbol{F}}_i^m + \overline{\boldsymbol{F}}_i^d, \tag{10}$$

where

$$\overline{\boldsymbol{F}}_i^d = \overline{\boldsymbol{\xi}}(t), \qquad \overline{\boldsymbol{F}}_i^m = \sum_{j\in\{\text{neigh.}\}} \overline{\boldsymbol{F}}_{ij}^m,$$
$$\overline{\boldsymbol{F}}_{ij}^m = \begin{cases} \overline{\mu}_s \left[\exp\left(-\overline{a}_s(\overline{\delta}_{ij}-2)\right) - \exp\left(-2\overline{a}_s(\overline{\delta}_{ij}-2)\right)\right] \frac{1}{\overline{\delta}_{ij}} \left(\overline{\boldsymbol{x}}_i - \overline{\boldsymbol{x}}_j\right), & \text{if } \overline{\delta}_{ij} \le 2, \\ \overline{\mu}_l(\text{FBS}_{ij}) \left[\exp\left(-\overline{a}_l(\overline{\delta}_{ij}-2)\right) - \exp\left(-2\overline{a}_l(\overline{\delta}_{ij}-2)\right)\right] \frac{1}{\overline{\delta}_{ij}} \left(\overline{\boldsymbol{x}}_i - \overline{\boldsymbol{x}}_j\right), & \text{otherwise.} \end{cases} \tag{11}$$

Here, we defined $\overline{\delta}_{ij} = \|\overline{\boldsymbol{x}}_i - \overline{\boldsymbol{x}}_j\|$, $\overline{\mu}_s = \mu_s/r$, $\overline{\mu}_l = \mu_l/r$, $\overline{a}_s = a_s r$, and $\overline{a}_l = a_l r$. The Gaussian noise, $\overline{\boldsymbol{\xi}}$, is characterised by zero mean, $\langle\overline{\boldsymbol{\xi}}_i(t)\rangle = 0$, and the covariance, $\langle\overline{\boldsymbol{\xi}}_i(t), \boldsymbol{\xi}_i(t')\rangle = 2\overline{D}_i\delta(t-t')$, with $\overline{D}_i = D_i/r^2$. From Eq (8), the rescaled cell motility coefficient is then given as follows

$$\overline{D}_i = \overline{D}(\text{FBS}_i) = \begin{cases} \overline{k} \cdot \text{FBS}_i + \overline{b}, & \text{if FBS}_i \le 5, \\ 1.7756, & \text{otherwise,} \end{cases} \tag{12}$$

with $\overline{k} = k/r^2$, $\overline{b} = b/r^2$ and $\text{FBS}_i = \text{FBS}(\overline{\boldsymbol{x}}_i)$ being the FBS level sensed by cell $i$. We also rescale Eq (9) describing the evolution of the FBS in culture medium:

$$\frac{\partial\text{FBS}}{\partial t} = \overline{D}_{fbs}\nabla^2\text{FBS} - \overline{k}_u\rho(\overline{\boldsymbol{x}})\text{FBS}. \tag{13}$$

In this equation, $\overline{D}_{fbs} = D_{fbs}/r^2$ and $\overline{k}_u = k_u/r^2$. In addition, the local cell density, $\rho(\overline{\boldsymbol{x}})$, is now calculated on the rescaled FEM elements, whose sides are scaled with $r$.

Parameter values after rescaling space with cell radius are listed in Extended Data Table 1 (lower part).

#### I.4 Metrics for aggregate quantification

We employ the DBSCAN (Density-Based Spatial Clustering of Applications with Noise) algorithm from Python Ester et al. (1996) to analyse cell aggregation patterns in our simulations. DBSCAN enables the detection of clusters by identifying densely packed points (cell centres in our case) based on the distances between them. The algorithm relies on two key parameters: $\epsilon$, which defines the maximum distance between two points to be considered neighbours, and *min_samples*, the minimum number of points required to form a cluster. For our analysis, we set $\epsilon = 2.1r$ and $min\_samples = 1$, ensuring that even small groups of cells are identified as clusters. Once the clusters are identified, we compute several metrics to quantify the average structure and composition of the aggregates:

- Average **cell number** in aggregates at a given time point. This metric is computed as the mean number of cells per aggregate across all aggregates at the time of interest, averaged over multiple numerical realisations.

- Average **aggregate area**, calculated as the total area covered by all cells within an aggregate. The area of an individual aggregate is determined by summing the areas of all pixels occupied by at least one cell, assuming a circular geometry with radius $r$ for each cell. This value is averaged over all aggregates at the given time point across multiple realisations.

- Average **aggregate density**, computed as the ratio of the average cell number to the average aggregate area.

#### I.5 Note on comparison of model simulations with experiments

Aggregate size and number vary due to stochasticity in motility and initial placement. We also note that, in experiments, cells experience a directed flux of liquid medium due to the non-flat bottom of the wells used for *Capsaspora* cultivation. This flow influences cell movement and can detach individual cells from the outer edges of multicellular aggregates. Since this effect is not intrinsic to the aggregation mechanism–*Capsaspora* does not aggregate without externally supplied FBS–we did not incorporate it in our simulations. Despite this discrepancy, our model effectively captures *Capsaspora* aggregation dynamics as can be appreciated in Figure 2c-d, Extended Data Figure 2, Movie 2 and Movie 3.

### II Supplementary movie captions

**Movie 1 Aggregation process of *Capsaspora owczarzaki* chemically induced with FBS.** Aggregation was induced by the addition of FBS (time $0$) to a final concentration 5% (v/v) in duplicates during different days. Aggregates were imaged every minute for $72$ h, and converted to a movie at $24$ fps. Time represents hh:mm. Scale bar represents $100$ $\mu m$. This movie can be accessed at https://doi.org/10.6084/m9.figshare.29044817.

**Movie 2 Comparison of experimental and simulated *Capsaspora* aggregation under 5% FBS.** Time-lapse animation of *Capsaspora* aggregation under 5% FBS. The right panel shows experimental recordings, while the left panel presents numerical simulations in a $1080 \times 1080$ $\mu m$ domain. In the simulation, cell colours indicate local FBS concentration, with depletion shifting from grey to dark blue. Adhesion follows a sigmoidal Hill response (see Supplementary Information). Parameter values are in Extended Data Table 1. This movie corresponds to Figure 2 of the main text.

**Movie 3 Comparison of experimental and simulated *Capsaspora* aggregation under 5% FBS in a small domain.** Time-lapse animation of *Capsaspora* aggregation under 5% FBS in a $360 \times 360$ $\mu m$ domain. The right panel shows experimental recordings, while the left panel presents numerical simulations. In simulations, cell colours indicate local FBS concentration, shifting from grey to dark blue as FBS depletes. The animation captures the full aggregation and disassembly process, corresponding to Extended Data Figure 2. Adhesion follows a sigmoidal Hill response (see Supplementary Information). Parameter values are in Extended Data Table 1.

**Movie 4 Comparison of experimental and simulated reaggregation in *Capsaspora udner 5% FBS.*** The Left panel shows experimental recordings of cell 24h after compleat dissociation, reaggregation is induced by the addition of FBS (new time $0$) to a final concentration 5% (v/v). Aggregates were imaged every minute. Time represents hh:mm.

Scale bar represents $100\ \mu m$. The top right panel presents numerical simulations in a $360 \times 360\ \mu m$ domain. cell colours indicate local FBS concentration, shifting from grey to dark blue as FBS depletes. Bottom right planel displays a zoomed $360 \times 360\ \mu m$ square marked in pink in the left panel for easier comparison with the model. Scale bar represents $100\ \mu m$.

**Movie 5 Reaggregation in *Capsaspora owczarzaki*.** Aggregation was induced by the addition of FBS to a final concentration 5% (v/v), after consumption of the FBS and dissociation of the aggregates FBS was readded in the same starting concentrations, generating a fast reaggregation response. Aggregates were imaged every minute. Time represents hh:mm. Scale bar represents $100\ \mu m$.

**Movie 6 Simulated *Capsaspora* aggregation under varying FBS levels for the Hill response model for cell adhesion.** Time-lapse animation of numerical simulations showing *Capsaspora* aggregation under different initial FBS concentrations, ranging from 0.5% (left) to 10% (right). In simulations, cell colours represent local FBS levels: red (high), grey (intermediate), and blue (low). The animation captures aggregation dynamics followed by disassembly upon FBS depletion. This video corresponds to Figure 3a. The numerical setup follows Figure 3a, with parameter values in Extended Data Table 1.

**Movie 7 Simulated *Capsaspora* aggregation under varying FBS levels for the moderate linear response model (shallow slope) for cell adhesion.** Time-lapse animation of numerical simulations showing *Capsaspora* aggregation under different initial FBS concentrations, ranging from 0.5% (left) to 10% (right). In simulations, cell colours represent local FBS levels: red (high), grey (intermediate), and blue (low). The animation captures aggregation dynamics followed by disassembly upon FBS depletion. This video corresponds to Figure 3b. The numerical setup follows Figure 3b, with parameter values in Extended Data Table 1.

**Movie 8 Simulated *Capsaspora* aggregation under varying FBS levels for the steep linear response model for cell adhesion.** Time-lapse animation of numerical simulations showing *Capsaspora* aggregation under different initial FBS concentrations, ranging from 0.5% (left) to 10% (right). In simulations, cell colours represent local FBS levels: red (high), grey (intermediate), and blue (low). The animation captures aggregation dynamics followed by disassembly upon FBS depletion. This video corresponds to Figure 3c. The numerical setup follows Figure 3c, with parameter values in Extended Data Table 1.

**Movie 9 Local cell density during *Capsaspora* aggregation and disassembly with the Hill response of cell adhesion to FBS.** Time-lapse animation of simulations illustrating local cell density during *Capsaspora* aggregation and disassembly with a sigmoidal Hill response to FBS. Cells are coloured by local density (upper colour bar), while the medium reflects FBS concentration (lower colour bar). Disassembly begins at the aggregate core around $55$ h, with the outer rim of cells persisting longer. Domain size: $360 \times 360\ \mu m$. Parameter values are in Extended Data Table 1. The same simulation, with cells coloured by local FBS levels, is shown in Movie 3.
